# FluxSprout: Single-cell metabolic modeling of vascular sprouting in zebrafish

**DOI:** 10.64898/2026.10.24.761567

**Authors:** Mohammad Reza Zargar, Ratul Chowdhury

## Abstract

Genome-scale metabolic modeling offers a powerful framework for identifying cell-state-specific metabolic vulnerabilities, yet its application to heterogeneous biological systems remains limited by uncertainty in network annotations, thermodynamic feasibility, transcriptome-to-flux inference, and target specificity. Here, we introduce FluxSprout, a single-cell transcriptomics-informed genome-scale metabolic modeling framework to resolve metabolic heterogeneity during zebrafish sprouting angiogenesis. By integrating single-cell RNA sequencing of 10,090 quality-controlled cells with Zebrafish-GEM, we reconstructed network ensembles for 14 endothelial and hemogenic states. We established four safeguards: stoichiometric auditing, loop auditing, cell-cycle-controlled testing, and ensemble specificity screening. First, we identified four inorganic pyrophosphatase reactions misannotated as oxidative phosphorylation, showing that apparent pathway essentiality arose only with unbounded exchange fluxes. Second, 19.5% of reactions could carry thermodynamically infeasible loop flux, including every candidate of a conventional screen. Third, tip-like and stalk-like cells differed mainly in cell-cycle phase, and elevated glycolysis separated sprouting from established vessels only in gene expression. In contrast, sprouting vessels had higher predicted growth capacity at both constraint strengths, with serine and nucleotide synthesis following in fixed proportion. Fourth, screening 825 genes across four network-extraction settings and 13 non-endothelial lineages left no candidate called by more than one setting. Notably, SLC25A20, that screen’s lead candidate, had no predicted effect in any FASTCORE network. Although the data derive from a single pooled clutch and require experimental validation, FluxSprout provides a transferable, uncertainty-aware framework for distinguishing genuine metabolic dependencies from artifacts of network reconstruction, model assumptions, and cellular heterogeneity, and identifies biosynthetic capacity as a testable feature of sprouting vessels.

## 1. Introduction

Sprouting angiogenesis, the process by which new vessels branch from existing ones, underlies development and wound healing and is dysregulated in diseases in which vessel growth is either pathologically excessive, as in tumor angiogenesis, or insufficient, as in ischemia [1], [2]. The process is coordinated by a division of labor between endothelial cell (EC) states: migratory, filopodia extending tip cells lead each sprout, proliferative stalk cells elongate it, and quiescent arterial and venous cells maintain the vessel wall behind it [3]. This specification is classically attributed to VEGF-A gradients acting through VEGFR2 and to the DLL4–Notch lateral inhibition circuit, which continuously reassigns the tip position among neighboring cells [4], [5]. Because the assignment is dynamic rather than fixed, the states are best understood as interconvertible phenotypes of one lineage, which is precisely what makes a description resolved by state rather than averaged over tissue necessary.

Over the past decade this signaling picture has been joined by a metabolic one. Tip cells were shown to rely disproportionately on glycolysis, driven by the phosphofructokinase-2/fructose-2,6-bisphosphatase isoform PFKFB3 [6], with pharmacological or genetic PFKFB3 suppression normalizing pathological vessel branching in vivo [7], [8] and with glycolytic flux governing the cell rearrangements that underlie tip–stalk shuffling [9]. Proliferating stalk cells were reported to depend on CPT1a-mediated fatty acid oxidation (FAO) to supply carbon for deoxyribonucleotide synthesis [10], whereas quiescent ECs upregulate FAO for redox homeostasis and vasculoprotection [11], and lymphatic ECs require the same pathway for lymphangiogenesis [12]. Serine and one-carbon metabolism via phosphoglycerate dehydrogenase [13], glutamine and asparagine metabolism [14], [15] and fatty acid synthase [16] have separately been implicated in EC proliferation and vessel formation, and the field has been synthesized into an account of endothelial identity centered on metabolism [17], [18]. The practical appeal of this account is that metabolic enzymes are druggable in ways that VEGF signaling has proven not to be, given the resistance and vascular toxicity that anti-VEGF agents encounter [19].

This account is not without internal tension. Independent functional evidence indicates that mitochondrial complex III, rather than glycolysis, is specifically required for EC proliferation during angiogenesis [20], direct measurements in tip-like cells assign roles to both glycolysis and respiration [21], and tip cells isolated in vitro were found to be less, not more, glycolytic than non-tip cells [22]. Much of the supporting evidence rests on bulk or pooled measurements and single gene perturbations rather than on single-cell, statistically controlled or flux level data. Single-cell atlases of murine and tumor endothelium have since established that EC states are transcriptionally heterogeneous and metabolically plastic [23], [24], [25], but they characterize expression rather than flux, and an expression difference need not imply a flux difference once stoichiometric and mass balance constraints are imposed across the network. A further caveat is that canonical mammalian tip markers do not transfer cleanly to zebrafish, in which *dll4*, *esm1* and *cxcr4a* mark arterial identity rather than tip identity in the early trunk, while *apln*, *aplnrb* and *marcksl1a* better capture the tip program [26], [27].

Proliferation is a second confound. Cell-cycle programs dominate the transcriptomes of growing tissue and can by themselves separate cells into clusters [28], and because tip and stalk cells differ partly in proliferative status, a boundary between them in single-cell data may reflect cell-cycle phase rather than position in the sprout; in the libraries analyzed here, the original study did not resolve tip and stalk cells at all [26]. Zebrafish is nonetheless an informative system in which to test these claims, because intersegmental vessel sprouting is stereotyped in position and timing, amenable to live imaging and genetically tractable [29], [30], [31], so that a computational prediction made in this system can in principle be falsified by a targeted perturbation in the same system.

Genome-scale metabolic models (GEMs) translate gene expression into predicted reaction flux under stoichiometric and mass balance constraints [32], [33], implemented in mature constraint-based toolboxes [34], [35], and variants informed by expression, such as E-Flux, scale reaction bounds continuously rather than binarizing the network [36]. Context-specific extraction algorithms prune a reference reconstruction to the reactions supported in a given sample [37], [38], [39], and their behavior depends strongly on thresholding and algorithmic choices [40], [41], [42]. A further source of error is thermodynamic: flux distributions can contain internal cycles that carry flux with no net conversion of substrate, and unless these are removed, they inflate flux variability ranges and the flux ratios computed from them [43], [44], [45]. The Human-GEM[46] family, including Zebrafish-GEM[47] derived from it by orthology, provides a shared namespace and curation standard across species, and network analyses of such models have been used to separate core from context-specific metabolism across human tumors [48]. Single-cell constraint-based modeling is an active area [49], [50], and related spatially resolved work has shown that flux-based prioritization can nominate transporter level vulnerabilities that expression analysis alone does not surface [45]; integrating such models with biomolecular simulation is a stated next step for the field [51].

No study, to our knowledge, has combined single-cell transcriptomic resolution with constraint-based flux modeling to characterize the metabolic states of sprouting angiogenesis, and the methodological question of how such a study should guard against its own false positives has received less attention still. Both gaps matter here for the same reason. Averaged over cell types, expression cannot distinguish a genuine per cell metabolic program from a marker gene artifact or from a difference in cell-cycle phase, and a pathway score based on expression cannot by itself establish whether a pathway is dispensable, essential or selectively required, because that determination depends on the redundancy of the surrounding network, on the reaction list assigned to the pathway and on the demands the objective function places on the cell. The aim of this study was therefore to reconstruct endothelial state-specific metabolic networks in the sprouting zebrafish trunk and, using them as a worked case, to develop and report four checks that a constraint-based vulnerability screen can be held to: an audit of reaction stoichiometry against the subsystem labels a pathway analysis relies on; an audit of thermodynamically infeasible cycles, which identifies the reactions whose flux ranges and ratios cannot be read without loop removal; a test at two levels in which each headline claim of the field is treated as a falsifiable hypothesis at both the marker and the flux level, with cell-cycle phase controlled and with metacells rather than resampled cells as the unit of replication; and a specificity control in which every candidate is evaluated across an ensemble of extraction methods and thresholds and in every non-endothelial lineage of the same dataset. Each check changed a conclusion we would otherwise have reported, and we present the angiogenesis results together with the checks that qualify them.

## 2. Materials and Methods

### 2.1. Single-cell RNA sequencing data and quality control

Single-cell RNA sequencing data of 24 hpf zebrafish embryos were obtained from Gurung *et al.* [26], who dissociated whole embryos from crosses of the etv2^ci32Gt^; *UAS:GFP* gene-trap line with *Tg(kdrl:mCherry)* and sorted, by fluorescence-activated cell sorting, a GFP⁺mCherry⁺ vascular endothelial fraction (GEO accession GSM6138416, 6,390 droplets) and a GFP⁺mCherry⁻ fraction of etv2 reporter-positive, kdrl-negative cells (GSM6138417, 6,990 droplets), which is dominated by erythroid and myeloid derivatives of etv2⁺ progenitors rather than being a general sample of non-vascular tissue, for 13,380 droplets in total prior to filtering (Table 1). Filtered feature barcode matrices were loaded per sample with Scanpy v1.12[52] and concatenated by outer join with zero filling, after per sample quality control metrics had been computed, so that mitochondrial and ribosomal transcript fractions were calculated before any between samples normalization could distort them.

**Table 1.** Single-cell RNA sequencing datasets and quality control. Embryos from *etv2*^ci32Gt^*; UAS:GFP* × *Tg(kdrl:mCherry)* crosses were sorted at 24 hpf into GFP⁺ mCherry⁺ and GFP⁺ mCherry⁻ fractions, and the two libraries were analyzed together. Raw droplets are counts before filtering; quality control-passing cells satisfy at least 500 detected genes, at least 1,000 unique molecular identifiers, fewer than 5% mitochondrial reads and doublet removal.

| GEO accession | Sort gate (24 hpf) | Raw droplets | QC-passing cells |
| --- | --- | --- | --- |
| GSM6138416 | GFP<sup>+</sup> mCherry<sup>+</sup> (*etv2* reporter<sup>+</sup>, *kdrl*<sup>+</sup>) | 6,390 | 5,066 |
| GSM6138417 | GFP<sup>+</sup> mCherry<sup>−</sup> (*etv2* reporter<sup>+</sup>, *kdrl*<sup>−</sup>) | 6,990 | 5,024 |
| Total | — | 13,380 | 10,090 |

Cells were retained if they had at least 500 detected genes, at least 1,000 total unique molecular identifiers and fewer than 5% mitochondrial reads. Upper tail total count outliers, defined as more than five median absolute deviations above the median on a log scale, were to be removed as likely multiplets (this filter removed no droplet in either library), as were droplets with a Scrublet doublet score above 0.25 computed independently per sample [53]; genes detected in fewer than three cells were discarded. This procedure yielded 10,090 quality-controlled cells, comprising 5,066 GFP⁺mCherry⁺ and 5,024 GFP⁺mCherry⁻ cells, across 24,205 genes, and removed 3,290 droplets in total, of which 2,256 failed the gene or transcript threshold, 1,921 the mitochondrial threshold and 128 the doublet threshold, with overlaps (Fig. 1D).

**Fig. 1.**
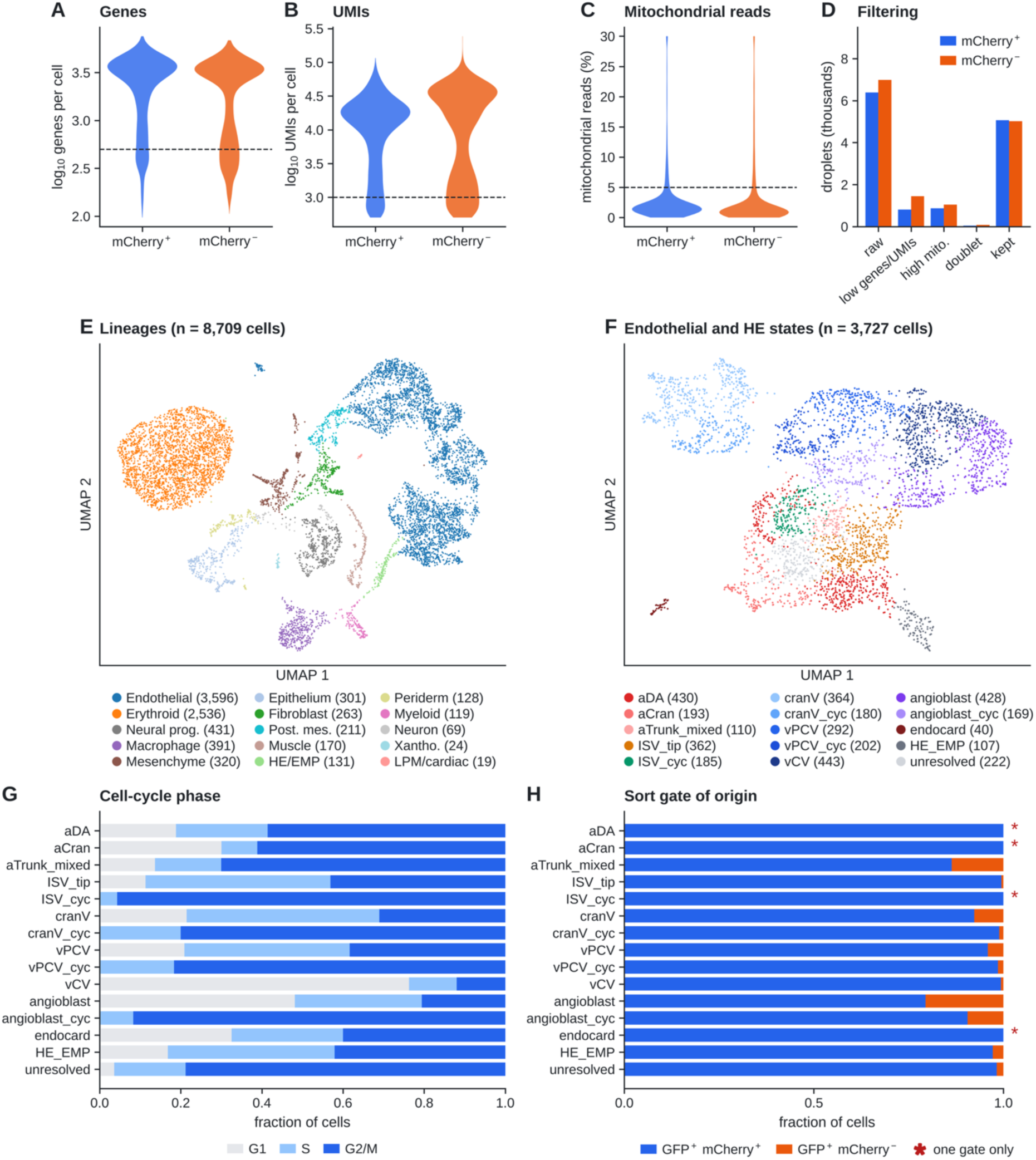
Quality control, lineage annotation and endothelial states of 24 hpf zebrafish single-cell libraries. Per sample distributions of genes per cell **(A)**, transcripts per cell **(B)** and mitochondrial read fraction **(C)**, with retention thresholds dashed; droplets removed at each quality control step, where the low-complexity bar is the union of the gene and transcript filters (**D)**. UMAP of the 8,709 annotated cells colored by lineage, with cell numbers **(E)**, and of the 3,727 endothelial and hemogenic cells colored by state **(F)**. Cell-cycle phase composition **(G)** and sort gate of origin **(H)** per state; “one gate only” marks states with no cell from the GFP⁺mCherry⁻ gate. The GFP⁺mCherry⁻ gate contains etv2 reporter-positive, kdrl-negative cells, not a general sample of non-vascular tissue.

### 2.2. Normalization, clustering and cell-type annotation

Counts were normalized to 10,000 reads per cell and log1p-transformed. Three thousand highly variable genes were selected with the Seurat v3 [54] flavor of the Scanpy implementation, batched by sample so that genes variable in only one sorted fraction were not preferentially selected, and transgene, mitochondrial, ribosomal and hemoglobin genes were excluded from the embedding to prevent the sort marker itself and dominant erythroid transcripts from driving the structure. The data were then scaled, and principal component analysis was computed on the top 50 components.

Harmony integration across the two sorted fractions [55] was computed and stored but deliberately not applied to the primary embedding. The two fractions are distinct sorted populations rather than technical batches of one population, so global correction would have removed genuine compositional differences; endothelial cells were drawn almost entirely from the GFP⁺mCherry⁺ gate (four of the fourteen endothelial and hemogenic states contain no cell from the other gate; Fig. 1H), so a correction across gates would have had little to align in the compartment of interest. Leiden clustering at resolution 1.0 [56] followed by annotation from markers resolved fifteen embryonic lineages. Four Leiden clusters (1,381 cells) were removed as low quality. They were identified by two joint criteria: a median of 998 detected genes and 3,377 transcripts per cell, against 3,406 and 21,405 in the retained clusters, and top marker sets composed of transgene, ribosomal, hemoglobin or apoptotic transcripts (eGFP with keratins; fabp3 with hmgb2b and hmgn2; ribosomal proteins with pmaip1; apoc1 and mfap4 with embryonic hemoglobins) rather than lineage-defining genes. These cells pass the per cell filters of Section 2.1 but cluster on RNA content rather than identity. This left 8,709 annotated cells. Cell-cycle phase was assigned from the S and G2/M gene sets of Tirosh *et al.* [57] mapped to zebrafish orthologs by name (including a and b paralogs; 45 S and 50 G2/M genes recovered). The 3,727 endothelial and hemogenic cells were then re-embedded (2,000 highly variable genes, 30 principal components) and subclustered at Leiden resolution 1.4 into 22 clusters, which were assigned to fourteen states from zebrafish vascular markers [26], [27] and from hox gene expression, which separates cranial (hox-negative) from trunk vessels. A further cluster of 222 cells with low complexity (median 1,278 genes) and ribosomal top markers was labeled unresolved and excluded from modeling. Cell labels are provided in the deposit, because Leiden numbering can change between software versions. Clustering stability was assessed by repeating the clustering on 80% and 90% random subsamples of the cells and quantifying agreement with the full solution by the adjusted Rand index (0.78 ± 0.03 over ten subsamples).

For statistical testing, the fourteen states were mapped onto four compartments defined by markers, and because three of the four pool more than one state we give the membership explicitly. **Arterial** (n = 623) pools the trunk dorsal aorta (aDA, 430) with the hox-negative cranial arterial cluster that contains the aortic arches (aCran, 193). **Venous** (n = 1,099) pools the posterior cardinal vein (vPCV, 292), caudal vein (vCV, 443) and cranial venous (cranV, 364) clusters and excludes their cycling counterparts. **Tip** (n = 362) is the intersegmental vessel cluster expressing esm1, hlx1, apln and aplnrb (ISV_tip). **Cycling endothelium** (n = 736) pools the cycling intersegmental vessel cluster (ISV_cyc, 185), which carries the same sprouting program as ISV_tip, with the cycling posterior cardinal vein (vPCV_cyc, 202), cranial venous (cranV_cyc, 180) and angioblast (angioblast_cyc, 169) clusters. Because ISV_tip and ISV_cyc share one program and differ mainly in cell-cycle phase (Section 3.2), we treat them together as the **sprouting** population (n = 547) in the flux and knockout analyses, and compare it with the **established** vessels (aDA, aCran, vPCV, vCV and cranV; n = 1,722). The endocardial, non-cycling angioblast, mixed trunk arterial and hemogenic clusters were retained in the atlas but excluded from these compartments, because the claims under test do not address their metabolic identity. Group membership is given in Table S3 and is used unchanged in every analysis.

### 2.3. Genome-scale metabolic model and curated pathway definitions

Zebrafish-GEM (12,833 reactions, 8,435 metabolites and 2,709 genes), a derivative of Human-GEM obtained by orthology mapping, was loaded with COBRApy [34]. Of the 2,709 model genes, 2,392 (88.3%) were present in the quality-controlled dataset, and the remainder were left at default bounds rather than being switched off, so that absence of detection in a sparse single-cell library was not silently treated as absence of the enzyme. Flux consistency was assessed with all exchange reactions open, which retained 11,356 reactions, and again under the embryonic medium described below, which retained 9,480 reactions and 2,170 genes; the latter network is the reference from which every model in this study was derived.

Ten core pathways were curated from the subsystem annotations of the reconstruction, each pathway being the union of one or more named subsystems so that the curation can be reproduced from the distributed model file: glycolysis and gluconeogenesis (43 reactions), the tricarboxylic acid (TCA) cycle (34), oxidative phosphorylation (10 by subsystem label, subsequently corrected to 6; Section 2.7), the pentose phosphate pathway (26), mitochondrial fatty acid oxidation (189, the union of the seven mitochondrial β-oxidation subsystems), fatty acid biosynthesis (166, the union of the four fatty acid biosynthesis subsystems), glutaminolysis (41, operationalized as the alanine, aspartate and glutamate metabolism subsystem), the mitochondrial carnitine shuttle (112), nucleotide synthesis (187, purine and pyrimidine metabolism, excluding the generic nucleotide metabolism subsystem) and serine and one-carbon metabolism (127, glycine–serine–threonine metabolism together with folate metabolism) (Table 2). The corrected oxidative phosphorylation set is used in every analysis reported here. Simulations were run on a yolk-like embryonic medium with uptake bounds of 10 for glucose, 20 for oxygen and 10 for glutamine, 1 for each of the other 19 amino acids, 1 for each of six fatty acids (palmitate, oleate, linoleate, linolenate, docosahexaenoate and arachidonate) and for cholesterol, choline and ethanolamine, 0.1 for each of fourteen vitamins and cofactors, and unbounded water, protons, phosphate, sulfate and iron; all secretions were open. Vitamin B12 (aquacob(III)alamin) and γ-tocopherol are included because, without them, the cofactor pool of the biomass reaction cannot be produced on a defined medium. The objective was the generic human biomass reaction MAR13082, which carries a growth-associated maintenance of 45 ATP; every biomass reaction is distributed with bounds of zero and was opened for this purpose, and the model’s default objective, MAR00021, which has no growth-associated ATP, was used only in the audit of Section 2.11. A non-growth-associated maintenance demand (NGAM) of 1 was imposed on the ATP hydrolysis reaction MAR03964. On this medium growth is set by essential amino acid supply (leucine has the only non-zero reduced cost at the optimum) while glucose and oxygen uptake are fully used, which we report because it determines how any change in carbon or oxygen supply can act on growth. Every exchange reaction and its bound, in the baseline medium and in each sensitivity medium, is listed in Table S2.

**Table 2.**
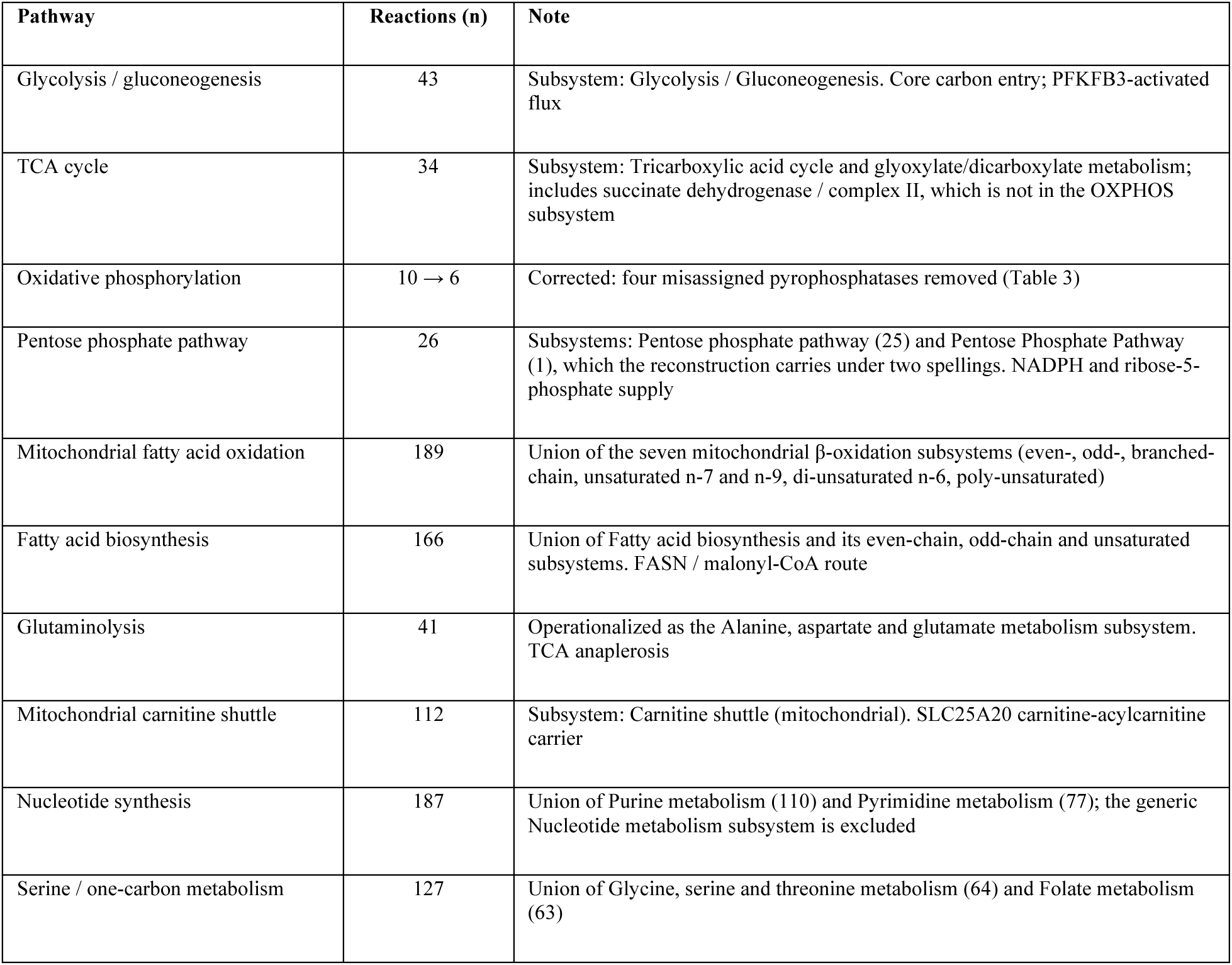
Curated metabolic pathway definitions. Ten core pathways curated from the subsystem annotations of Zebrafish-GEM. Each pathway is the union of one or more named subsystems, given in the Note column, so that the curation can be reproduced from the distributed model file; the reaction counts were verified against it. Oxidative phosphorylation was corrected from ten to six reactions after four inorganic pyrophosphatase reactions were found to be misassigned to the subsystem (Section 2.7 and Table 3).

| Pathway | Reactions (n) | Note |
| --- | --- | --- |
| Glycolysis / gluconeogenesis | 43 | Subsystem: Glycolysis / Gluconeogenesis. Core carbon entry; PFKFB3-activated flux |
| TCA cycle | 34 | Subsystem: Tricarboxylic acid cycle and glyoxylate/dicarboxylate metabolism; includes succinate dehydrogenase / complex II, which is not in the OXPHOS subsystem |
| Oxidative phosphorylation | 10 → 6 | Corrected: four misassigned pyrophosphatases removed (Table 3) |
| Pentose phosphate pathway | 26 | Subsystems: Pentose phosphate pathway (25) and Pentose Phosphate Pathway (1), which the reconstruction carries under two spellings. NADPH and ribose-5-phosphate supply |
| Mitochondrial fatty acid oxidation | 189 | Union of the seven mitochondrial β-oxidation subsystems (even-, odd-, branched-chain, unsaturated n-7 and n-9, di-unsaturated n-6, poly-unsaturated) |
| Fatty acid biosynthesis | 166 | Union of Fatty acid biosynthesis and its even-chain, odd-chain and unsaturated subsystems. FASN / malonyl-CoA route |
| Glutaminolysis | 41 | Operationalized as the Alanine, aspartate and glutamate metabolism subsystem. TCA anaplerosis |
| Mitochondrial carnitine shuttle | 112 | Subsystem: Carnitine shuttle (mitochondrial). SLC25A20 carnitine-acylcarnitine carrier |
| Nucleotide synthesis | 187 | Union of Purine metabolism (110) and Pyrimidine metabolism (77); the generic Nucleotide metabolism subsystem is excluded |
| Serine / one-carbon metabolism | 127 | Union of Glycine, serine and threonine metabolism (64) and Folate metabolism (63) |

### 2.4. GPR-aware E-Flux constraint scaling

Reaction level expression scores were computed by evaluating the gene–protein–reaction (GPR) Boolean rule of each reaction over mean per state expression. For a reaction *j* with gene set *G*ⱼ, AND relationships, which represent the obligate subunits of an enzyme complex, were mapped to the minimum and OR relationships, which represent interchangeable isozymes, to the maximum constituent gene value:

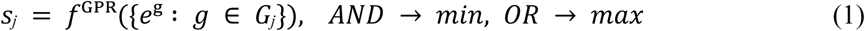

where *e*ᵍ is the mean normalized expression of gene *g* in the state under consideration. Genes absent from the dataset are ignored inside the rule, so that an undetected gene neither closes nor opens a reaction; a reaction none of whose genes is detected keeps its default bounds.

Scores were normalized to their 95th percentile across reactions, so that no single highly expressed reaction sets the scale for the whole network, and the bounds of each reaction were scaled by a capacity U, following the E-Flux principle of continuous rather than binary constraint [36]:

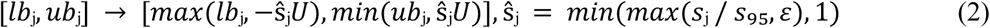

with a floor *ε* = 0.05. The capacity matters because the scaled bound must be comparable to the fluxes the medium allows for expression to constrain anything. When bounds are scaled from the default of ±1000 (U = 1000), the tightest expression bound (50) is several times larger than the glucose uptake of 10, and in our models a median of only 8 reactions reached their expression bound, with 87% of the unconstrained growth retained. We therefore swept U from 1000 to 20 (Fig. 5A) and used U = 50 as the primary setting, at which the median state model retains about half of reference growth and 33 reactions reach their expression bound, with U = 100 as a sensitivity setting; the conventional scaling (normalization to the maximum, U = 1000) is reported alongside for comparison. Both bounds of a reversible reaction are scaled by the same factor, so scaling narrows a reversible range symmetrically and does not shift its direction. We used the GPR rule rather than a flat mean of the associated genes, because a flat mean understates flux through OR-linked reactions with multiple isozymes; the most consequential case was PFKFB3, whose regulatory role is distributed across several paralogs in zebrafish. Exchange reactions, the biomass reaction, the four biomass pool reactions and the maintenance reaction were exempted from scaling.

### 2.5. Flux analysis: pFBA, loop-corrected FVA and metacell models

Growth-optimal flux distributions were obtained by parsimonious flux balance analysis, which selects among alternative optima the distribution of minimum total absolute flux [58]. Because minimizing total flux also removes internal cycles, every pFBA solution was checked against its nearest loop-free flux distribution by CycleFreeFlux [44]; the largest L1 distance across all models was below 10⁻⁷, so the pFBA solutions reported here are loop-free. Feasible ranges were obtained by flux variability analysis at 90% of optimal growth [59], both as computed and after loop correction: each extreme was tested against its nearest loop-free flux distribution and, where it did not survive, the reaction was re-optimized within the sign pattern of that loop-free distribution. The corrected range is a conservative estimate of the loop-free range, and reactions whose range changed are flagged as loop-dependent (Table 6).

Rather than resampling cells to recompute a single state profile, which barely moves a mean taken over hundreds of cells, we used metacells as the unit of replication. Within each state and lineage, cells were partitioned by k-means on the first 30 principal components into groups of about 20 (*k* = *max*(3, *n*/20); groups below 5 cells discarded) [60], giving 384 metacells; E-Flux models were built for the 325 in the endothelial and hemogenic states and five non-endothelial lineages. Flux phenotypes were summarized as the glycolytic share of ATP production (phosphoglycerate kinase plus pyruvate kinase over these plus ATP synthase), lactate secretion per unit glucose uptake, oxygen uptake, phosphoglycerate dehydrogenase (PHGDH) flux, carnitine–acylcarnitine carrier flux, and de novo purine and pyrimidine synthesis per unit growth (PPAT and CAD fluxes divided by growth). The last two are reported because, in a growth-maximizing model, biosynthetic fluxes are fixed by the biomass composition and therefore cannot rank cell states by anything other than growth (Section 3.6).

### 2.6. Context-specific model ensemble

As a complement to the expression-scaled models, in which the network is bound constrained but not pruned, state-specific models were extracted from the 9,480-reaction reference network for each of the fourteen endothelial and hemogenic states and each of the thirteen non-endothelial lineages (27 groups). Because the outcome of extraction is known to depend on both the algorithm and the threshold [40], [41], [42], we built an ensemble of four model sets rather than a single one. Three used FASTCORE [37] with core reactions defined by local-T2 thresholding [42] on log-transformed mean expression, with the global lower and upper thresholds set at the 15th and 65th (loose), 25th and 75th (default) and 40th and 85th (strict) percentiles of all gene–group values; a gene between the two thresholds is called active in a group if it is expressed above its own mean across groups. The fourth used GIMME [39], with the default local-T2 upper threshold as the expression cutoff and 90% of optimal growth as the required function.

Two practical points affect whether extracted models are usable and are reported for reproducibility. FASTCORE’s linear programs require every core reaction to carry flux above a small threshold, and with a maintenance demand imposed this cannot be satisfied when exchange reactions are themselves forced into the core; extraction was therefore run with the maintenance demand released and with exchanges outside the core. In addition, the support threshold drops micronutrient routes whose fluxes are about a millionth of the biomass flux, which leaves the biomass reaction unable to carry flux; after extraction we therefore added the minimal-flux set of reference reactions needed for biomass at 10% of the reference optimum with the maintenance demand restored. FASTCORE models contained 6,432– 7,626 (loose), 6,076–7,551 (default) and 5,414–7,218 (strict) reactions, and GIMME models 2,695– 3,905. Agreement between reaction sets, measured by the mean Jaccard index per group, was 0.94 between the loose and default FASTCORE sets, 0.91 between the strict and default sets, and 0.48 between FASTCORE and GIMME, so about half of an extracted network depends on the choice of algorithm.

### 2.7. Oxidative phosphorylation reaction audit

In the conventional screen (Supplementary Note S1), titrating the ten reactions labeled oxidative phosphorylation, with every exchange open, suggested that about 64% of growth depended on the pathway. Because this figure was large, and because a clean result of that kind is a reason for more scrutiny rather than less, the ten reactions were audited individually against the stoichiometric equations, compartment assignments and gene–protein–reaction rules stored in the reconstruction rather than against their subsystem labels. Four reactions (MAR03975, MAR03977, MAR03979 and MAR06912), all named “diphosphate phosphohydrolase” and encoding inorganic pyrophosphatase activity (H₂O + PPi → H⁺ + 2 Pi) in the endoplasmic reticulum, cytosol, peroxisome and mitochondrion respectively, were reclassified as non-OXPHOS. All four share the gene–protein–reaction rule *lhpp or ppa1b or ppa2 or prune*, none of which encodes an electron transport or ATP synthase subunit. The remaining six were retained as the corrected set: complex I (MAR06921), complex III (MAR06918), complex IV (MAR06914), ATP synthase (MAR06916), an FAD-linked dehydrogenase entry reaction into the quinone pool (MAR06911, FADH₂ + ubiquinone ⇌ FAD + ubiquinol) and an alternative, superoxide generating cytochrome *c* oxidase variant (MAR13081), which is constrained to zero flux in the distributed reconstruction and therefore contributes nothing under either definition. The complete audit, with the stored equation, compartment, gene association and verdict for each of the ten reactions, is given in Table 3. Two further observations from the audit are reported: the reconstruction contains a fifth inorganic pyrophosphatase, MAR07794 in the nucleus, which carries the subsystem label “Miscellaneous” rather than “Oxidative phosphorylation”; and succinate dehydrogenase (MAR04652) is assigned to the TCA cycle subsystem, not to OXPHOS, so complex II is outside this reaction set under both definitions. All analyses that depend on OXPHOS were rerun under both the legacy and the corrected definitions, together with blocks of the four mislabeled pyrophosphatases alone and of all five (Section 2.11), and the corrected definition is reported as primary throughout.

**Table 3.**
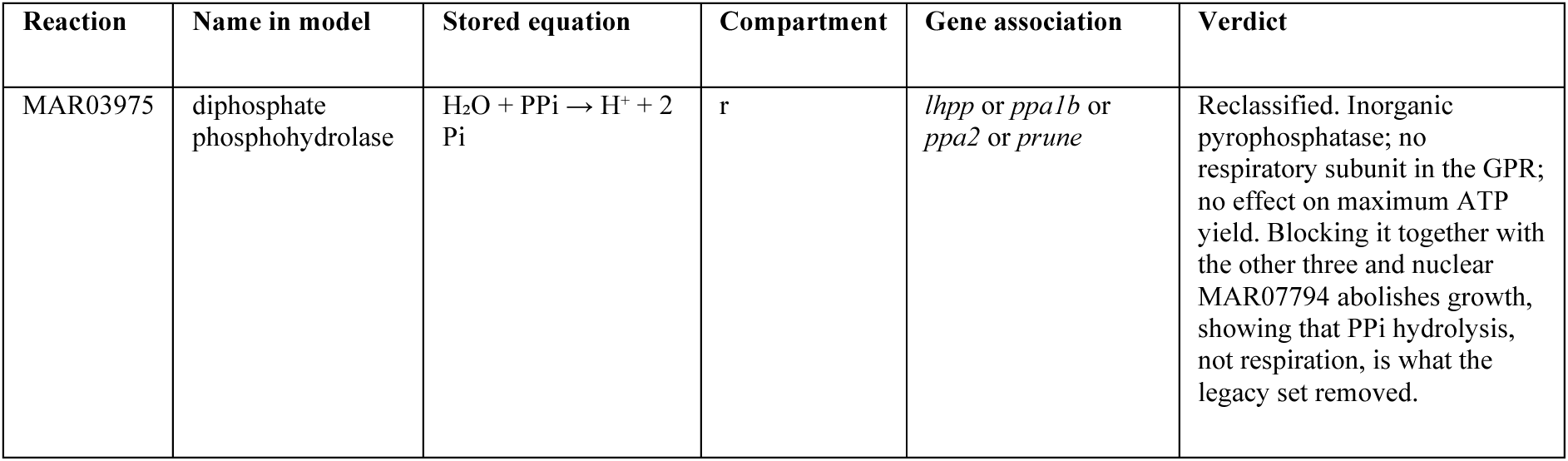
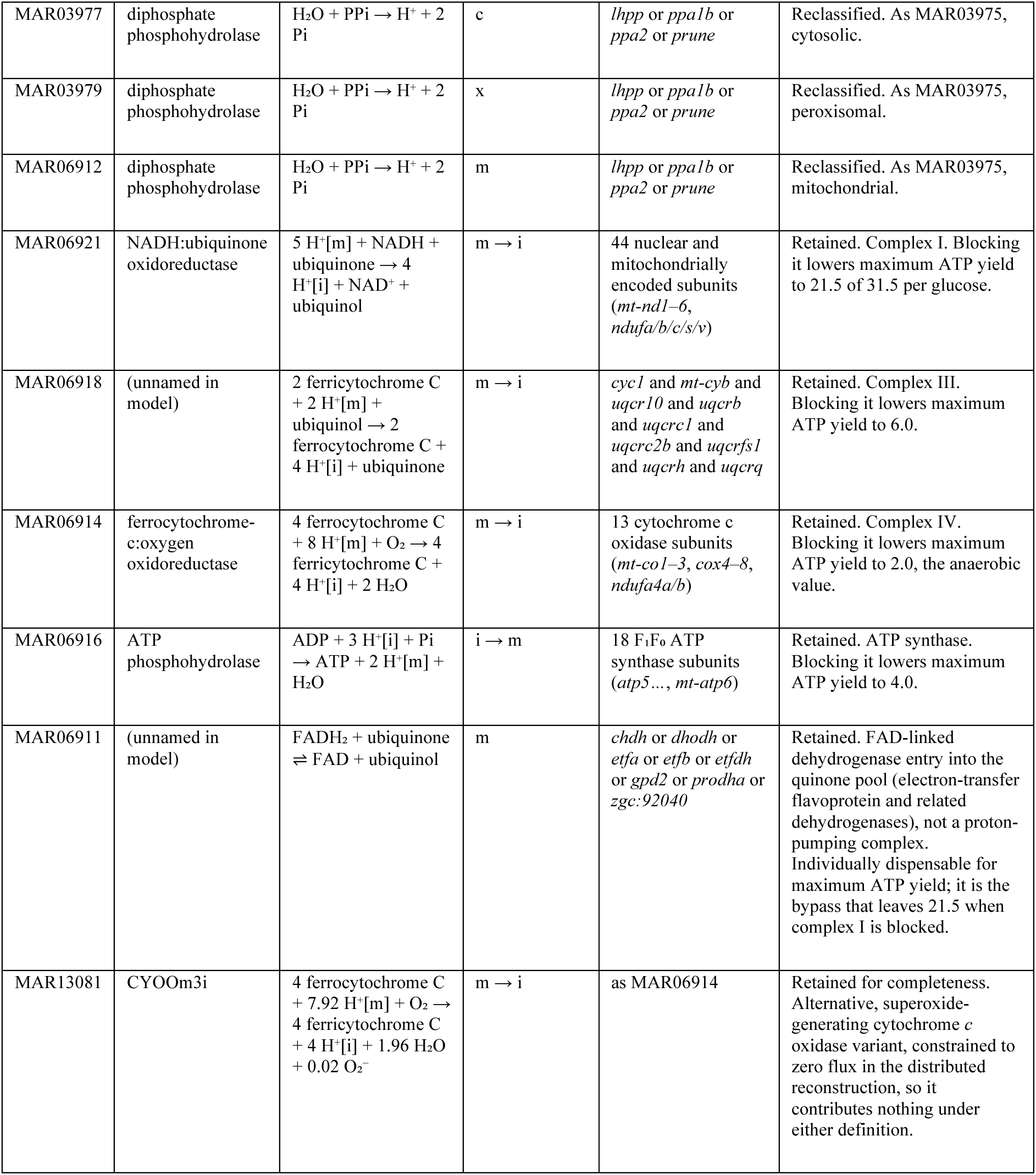
Reaction by reaction audit of the oxidative phosphorylation subsystem of Zebrafish-GEM. All ten reactions carrying the “Oxidative phosphorylation” subsystem label in the reconstruction, with the reaction name, stoichiometric equation and compartment as stored in the model file, the gene–protein–reaction rule, and the verdict of the audit. Four reactions encode inorganic pyrophosphatase activity and are reclassified; the six retained reactions form the corrected set used throughout. A fifth, identical pyrophosphatase reaction (MAR07794, nucleus) carries no OXPHOS label; blocking all five abolishes growth, whereas blocking only the four mislabeled ones does not. Every entry is taken directly from the distributed model file, so each reclassification can be checked from the stoichiometry and gene association alone. Compartments: c, cytosol; i, inner mitochondrion; m, mitochondrion; r, endoplasmic reticulum; x, peroxisome.

### 2.8. Audit of thermodynamically infeasible cycles

A reaction that can carry flux when the network exchanges no mass with its surroundings can do so only as part of an internal cycle, which by the loop law carries no thermodynamic driving force [43]. To find every such reaction we closed all exchange reactions, the biomass reactions and the maintenance reaction in the flux-consistent network and identified the reactions that could still carry flux, first by batched linear programs that maximize the number of reactions carrying a flux of at least 10⁻⁵ (in the manner of FASTCC [37]) and then by random-objective probes until five successive probes found no new reaction. Each gene was then characterized by the fraction of its GPR-associated reactions that are loop-capable. These reactions are not removed from the models, because loops do not by themselves change growth or knockout outcomes, but any flux range, flux ratio or sampled vertex involving them is reported only after loop correction (Section 2.5).

### 2.9. Statistical validation at the marker and flux levels

Per cell marker panel scores were computed with the Scanpy *score_genes* routine over gene panels built from the genes of each pathway’s Zebrafish-GEM reactions (the four pyrophosphatase reactions excluded) and detected in at least 5% of endothelial cells. Scores were compared between the focal compartment and the pooled remainder using three complementary procedures: bootstrap confidence intervals over 2,000 resamples, obtained by resampling both groups independently and recomputing the difference of means; a one-sided Mann–Whitney *U* test; and, because proliferation and sequencing depth both vary between compartments, an ordinary least squares model of the score on compartment membership with S score, G2/M score and log₁₀ transcript count as covariates, with heteroscedasticity-consistent standard errors. A claim was recorded as supported only if it was significant after correction and the focal compartment was also the highest scoring of the four. This rule was applied to the raw scores and, for the adjusted test, to the scores after removing the covariates. A claim whose confidence interval lay entirely on the opposite side of zero was recorded as opposite in direction rather than merely unsupported. False-discovery-rate correction was applied across the eight tested claims by the Benjamini–Hochberg procedure [61]. The same adjusted model was used to compare the two intersegmental vessel states, ISV_cyc against ISV_tip, for each of the eight pathway panels, with correction across panels.

Because marker level significance establishes only that an expression difference is real, each claim was also tested at the flux level, using the metacell models of Section 2.5 as replicates. Compartments were compared by two-sided Mann–Whitney tests, with Cliff’s delta as the effect size [62], and the Benjamini–Hochberg procedure was applied across all flux level tests (two capacities, five comparisons and eight read-outs). A flux level difference is described as robust only if it is significant at both U = 50 and U = 100 with the same sign.

Both levels of testing use the same 10,090 cells from a single pooled clutch. They therefore quantify within sample separation and its stability across metacells, and neither can estimate reproducibility across biological replicates; no test reported in this study should be read as evidence that a difference would recur in an independently raised clutch.

### 2.10. Vulnerability screening, specificity controls and human orthology

To show what each check changes, we also ran the screen as such screens are commonly performed (the conventional screen; Supplementary Note S1): oxidative phosphorylation defined by subsystem label, no loop correction, E-Flux bounds scaled from ±1000, a single FASTCORE extraction, and a specificity control against the erythroid and muscle models only. Its candidates (SLC25A20, SLC25A36 and IDH2) are carried through the analyses below as a reference. Every gene of the reference network whose deletion disables at least one reaction under its GPR rule (825 genes; the others are buffered by isozymes) was deleted individually in every model of the four-member ensemble, and growth after deletion was expressed as a fraction of growth before it. Inhibition in the sprouting population is the mean over ISV_tip and ISV_cyc, and within-endothelium selectivity is the difference between that and the mean inhibition across the established vessels. We state two calling rules before the results and apply both. The first, the two-lineage rule, calls a gene if it inhibits sprouting growth by at least 0.10 while leaving the erythroid and skeletal muscle models at full growth. The second, the all-lineage rule, which we treat as primary, additionally requires that the gene leaves all thirteen non-endothelial lineages at full growth and has a within-endothelium selectivity of at least 0.10. A gene is reported as a robust candidate only if it is called in more than one of the four ensemble members. For every gene called under the all-lineage rule, the cause of the call was traced by comparing the model in which the knockout is lethal with the closest model in which it is not (the other sprouting state where possible): with the knockout imposed in the union of the two networks, we found the minimal-flux set of reactions, present only in the unaffected model, that restores growth to at least half of its value.

Context sensitivity was assessed by reparametrizing the medium for three disease microenvironments and two further nutrient conditions, changing only the exchange bounds named in each case and leaving all others at their baseline values (glucose 10, oxygen 20, glutamine 10): tumor hypoxia (glucose 30, oxygen 1, glutamine 20); hyperglycemia (glucose 40, oxygen unchanged); and ischemia (glucose 2, oxygen 0.8, glutamine 2); carnitine supplementation (1), to test whether any carnitine-shuttle dependence reflects carnitine supply; and a tenfold reduction in every amino acid (0.1), because amino acids set growth on the baseline medium. Because flux balance analysis cannot lower maximum growth when a substrate bound is only relaxed, the hyperglycemia condition verifies that the medium is applied as intended and that the solver returns a monotone response; it is an implementation check and carries no biological information, and we do not count it as a validation. Recapitulation of published phenotypes (Section 3.8) was scored under a rule stated in advance: a phenotype counts as recapitulated if predicted flux through the target is elevated in the sprouting compartment relative to the quiescent one, that is, a ratio above 1.00, or if deleting the target reduces sprouting growth to 0.90 or below. Flux ratios were computed from loop-corrected pFBA solutions of pooled sprouting and established-vessel E-Flux models (U = 50), as the sum of absolute flux over the target gene’s reactions, and growth limitation was counted across the four ensemble members. Rows meeting the rule only through unconditional lethality are marked as such, since a gene that is lethal in every lineage is a general biosynthetic requirement rather than an endothelial phenotype. Candidate zebrafish genes were mapped to human orthologs through the DIOPT ortholog table shipped with Zebrafish-GEM [63], no DIOPT threshold was imposed; the support score is reported per gene and ranges from 3 to 10 across the mapped candidates, so the orthology of individual candidates varies in confidence and IDH2, at 3 of 10, is weakly supported, and the presence and reaction level conservation of each target was confirmed in Human-GEM. Software is described in Section 2.12.

### 2.11. Objective, maintenance demand, medium and flux caps

Because an apparent pathway dependence is a statement about the reaction set, the medium and the energy demand together, we repeated the oxidative phosphorylation titration as a full factorial on the flux-consistent reference network: two objectives (MAR13082, with 45 ATP of growth-associated maintenance, and MAR00021, with none), two media (the constrained embryonic medium, and all exchanges open), five NGAM values (0, 1, 5, 20 and 40) and four reaction sets (the legacy ten reactions, the corrected six, the four mislabeled pyrophosphatases, and all five pyrophosphatases including the nuclear MAR07794). Capacity was titrated from intact to fully blocked relative to each reaction’s flux in the intact optimum. Maximum ATP yield per glucose was computed by maximizing ATP hydrolysis on a medium of glucose (1), oxygen and inorganic ions, intact and with each reaction set or respiratory complex blocked. Finally, because a network with all exchanges open is limited only by the arbitrary bounds placed on internal reactions, we recomputed the open-medium optimum with those bounds set from 250 to 100,000 and counted the internal reactions sitting at the bound.

### 2.12. Implementation and reproducibility

Single-cell analyses used Scanpy [52]. Constraint-based analyses used COBRApy [34] for model handling and a purpose-written interface to the HiGHS linear programming solver [64] for FBA, pFBA, FVA, loop correction, gene deletion, FASTCORE and GIMME, which allowed the full knockout screen (825 genes, 27 groups and four ensemble members) to be run; with warm-started solves, a deletion takes about 0.03 s on the reference network. This interface reproduces COBRApy results on the E. coli core model to within 10⁻¹¹ for FBA, FVA and single-gene deletion, and finds the same blocked reactions. All analyses, with random seeds fixed, are contained in a single Jupyter notebook that regenerates every figure and supplementary table from the two GEO matrices and the distributed Zebrafish-GEM.

## 3. Results

### 3.1. An atlas of endothelial states at the onset of intersegmental vessel sprouting

To resolve the metabolic network expressed in each endothelial state at the onset of sprouting, we analyzed single-cell RNA sequencing data from 24 hpf zebrafish embryos in which a GFP⁺mCherry⁺ and a GFP⁺mCherry⁻ fraction had been sorted separately (Table 1). Of 13,380 droplets, 10,090 cells passed quality control (Fig. 1A–F). The two libraries behaved comparably, with median complexities of approximately 3.5 log₁₀ genes per cell in both fractions. The filters flagged 2,256 low complexity droplets, 1,921 with excessive mitochondrial content and 128 doublets; these sets overlap, and 3,290 droplets were removed in total, a loss dominated by poor-quality droplets rather than by doublets, as expected for whole embryo dissociations. Of the retained cells, 8,709 were confidently annotated into fifteen embryonic lineages spanning endothelial, hemogenic, erythroid, myeloid, macrophage, neural, muscle, mesenchymal and epithelial identities (Fig. 1E), with endothelium (3,596 cells) and erythroid cells (2,536) the most abundant.

Subclustering the 3,727 endothelial and hemogenic cells resolved fourteen states (Fig. 1F; markers in Fig. S1). These comprised a trunk dorsal aorta (aDA) and a hox-negative cranial arterial population that includes the aortic arches (aCran), a mixed trunk arterial cluster, posterior cardinal, caudal and cranial venous populations with cycling counterparts for the posterior cardinal and cranial veins, angioblast and cycling angioblast populations, an endocardial population, a hemogenic endothelium/erythromyeloid progenitor cluster marked by myb and runx1, and two intersegmental vessel (ISV) populations that both express the sprouting markers esm1, hlx1, apln and aplnrb. The two ISV populations differed sharply in cell-cycle composition: 96% of ISV_cyc cells were assigned to G2/M and none to G1, while ISV_tip contained 46% S-phase and 11% G1 cells (Fig. 1G). Endothelial cells were drawn almost entirely from the GFP⁺mCherry⁺ gate, and four states (aDA, aCran, ISV_cyc and endocardial) contain no cell from the other gate (Fig. 1H). The clustering was stable to subsampling, with an adjusted Rand index of 0.78 ± 0.03 between the full and 80–90% subsampled solutions.

### 3.2. Tip and cycling intersegmental vessel cells share one sprouting program and differ mainly in cell-cycle phase

Because tip and stalk identities in the sprout differ partly in proliferative status, we asked whether the two ISV populations differ in anything beyond cell-cycle phase before treating them as tip and stalk cells. Of the genes up in ISV_cyc relative to ISV_tip (449 at q < 0.05 and |log₂ fold change| > 0.5), 90% of the top 200 were correlated with the S or G2/M score across all endothelial cells (|r| > 0.3), and the 72 genes down in ISV_cyc were dominated by S-phase replication genes (rfc4, mcm7, pold1, fen1, gins2). When the S and G2/M scores were regressed out, the separation of the two populations fell from a silhouette of 0.18 to 0.06, and reclustering did not recover them (adjusted Rand index 0.12–0.31; Fig. 2A–C). Pathway marker scores told the same story: once phase and depth were controlled, ISV_cyc and ISV_tip did not differ in glycolysis (difference −0.008, q = 0.69), TCA cycle, fatty acid oxidation, serine and one-carbon, carnitine shuttle or pentose phosphate scores, and differed only in a lower oxidative phosphorylation score (−0.048, q = 0.03) and a higher nucleotide synthesis score (+0.019, q = 0.009) in ISV_cyc (Fig. 2D). In these data, therefore, the two ISV populations are best read as one sprouting population sampled in S and in G2/M, rather than as tip and stalk cells, consistent with the original study’s report that tip and stalk cells were not resolved [26]. In what follows we keep ISV_tip as a compartment for comparability with the literature, but test tip-versus-stalk claims within the ISV population, and pool ISV_tip and ISV_cyc as the sprouting population for flux and knockout analyses.

**Fig. 2.**
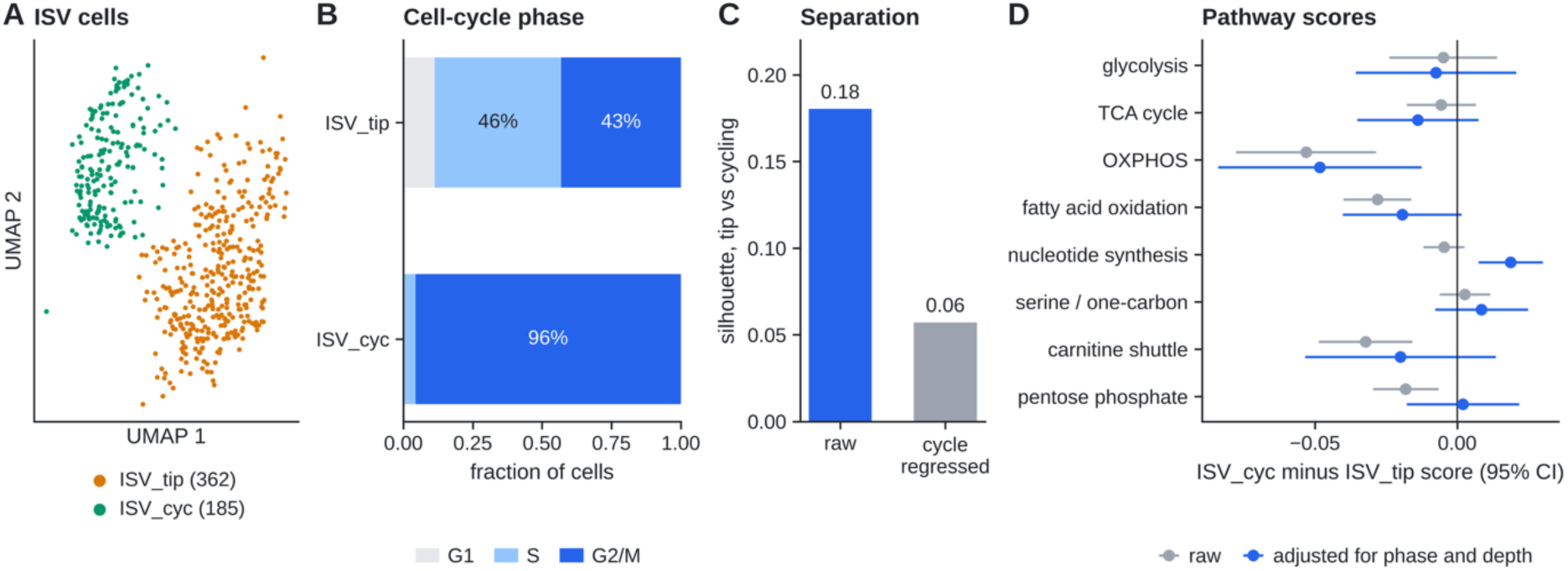
Tip and cycling intersegmental vessel cells differ mainly in cell-cycle phase. **(A)** The two ISV populations on the endothelial embedding. **(B)** Cell-cycle phase composition. **(C)** Separation of the two populations (silhouette on the first 20 principal components of non-cycle highly variable genes), before and after regressing out S and G2/M scores. **(D)** Difference in pathway marker score between ISV_cyc and ISV_tip, raw and adjusted for S score, G2/M score and sequencing depth (95% confidence intervals).

### 3.3. Marker level tests of the field’s metabolic claims

We next treated each headline claim of the field as a falsifiable hypothesis and tested it at two levels of stringency (Table 5). All per cell statistics below are computed on 10,090 cells from a single pooled clutch sorted into two libraries; they quantify separation within that sample and not reproducibility across clutches, and the *p*-values should be read accordingly. At the per cell marker level, five claims were supported by the raw (unadjusted) procedure: tip cell glycolytic elevation (Mann–Whitney p = 2.0 × 10⁻¹⁰), nucleotide synthesis elevation in the cycling endothelial compartment (p = 8.5 × 10⁻³²), serine/one-carbon elevation in the same compartment (p = 3.0 × 10⁻⁶), venous fatty acid oxidation elevation (p = 2.5 × 10⁻⁴) and tip cell oxidative phosphorylation (p = 1.3 × 10⁻¹⁵), all significant after Benjamini–Hochberg correction across the eight tested claims (Fig. 3A, Table 5). Because proliferation and sequencing depth both differ between compartments, each claim was retested with S score, G2/M score and depth as covariates (Fig. 3B). Four of the five survived; venous fatty acid oxidation did not (adjusted difference 0.003, q = 0.18), which shows that its raw signal is carried by the larger share of G1 cells in the venous compartment. The tip oxidative phosphorylation result survives with the corrected gene panel, from which the four pyrophosphatase genes are excluded; the misannotation of Section 3.4 affects reaction-set titration, not gene panel scores. Two claims were not supported. The proposal that the arterial lineage as a whole is the most glycolytic population failed under both procedures; the highest glycolytic score belonged to the tip state, which locates the effect at the sprouting vessel rather than at the whole lineage, and illustrates how a lineage level average can invert a state-level effect. The proposal that the cycling compartment carries the highest TCA cycle marker score also failed, the one-sided test in the proposed direction returning p = 0.98. Finally, the carnitine-shuttle score was lower, not higher, in cycling endothelium than in the rest (adjusted difference −0.015, confidence interval entirely below zero), which bears directly on any carnitine-shuttle vulnerability proposed for proliferating endothelium. Read together with Section 3.2, the tip glycolysis result is a difference between the sprouting ISV population and established arterial and venous vessels: within the ISV population, tip and cycling cells do not differ in glycolytic score.

**Fig. 3.**
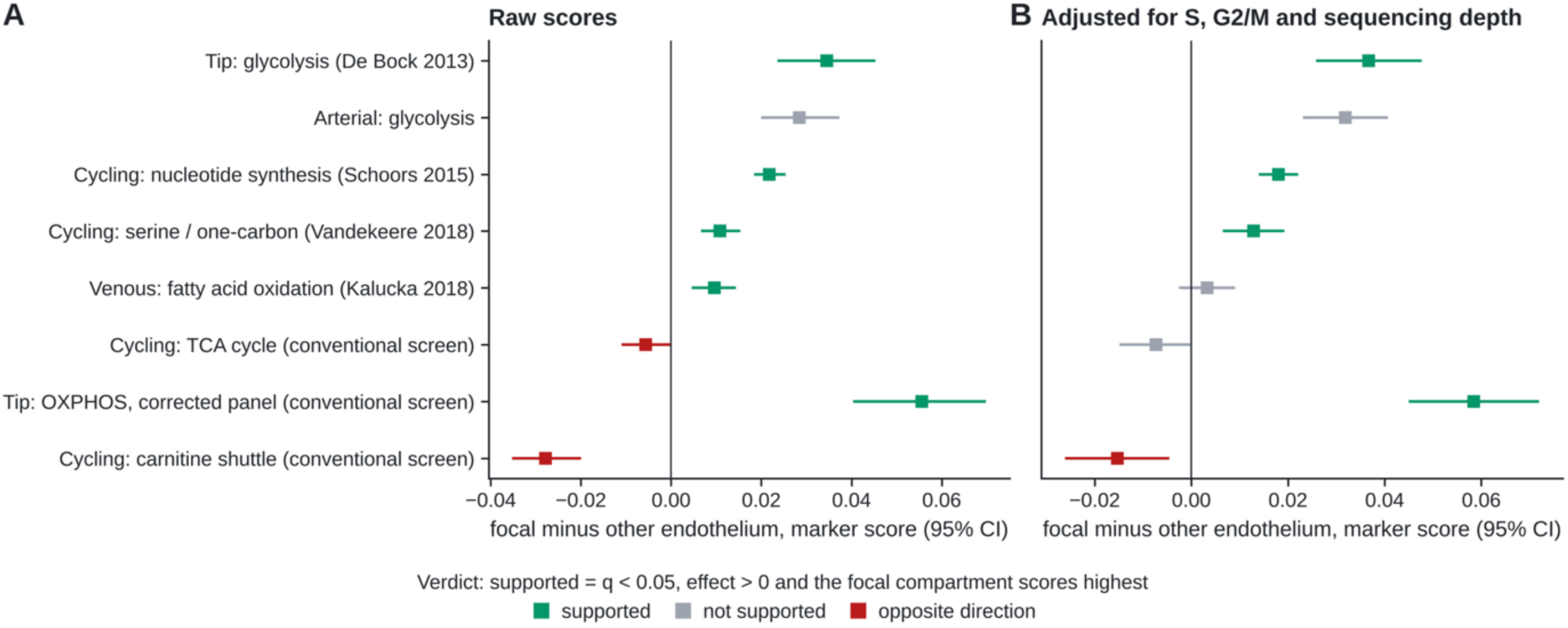
Marker level tests of eight metabolic claims. Difference in per cell marker score between the focal compartment and the pooled remainder, **(A)** raw, with 95% bootstrap confidence intervals, and **(B)** adjusted for S score, G2/M score and sequencing depth, with 95% confidence intervals from a heteroscedasticity-consistent linear model. Colors give the verdict recorded in Table 5: supported means q < 0.05, effect above zero and the focal compartment scoring highest of the four (the same rule in both panels).

### 3.4. Four pyrophosphatases are misannotated as oxidative phosphorylation, and apparent dependence on the pathway is set by objective and maintenance

In the conventional screen (Supplementary Note S1), titrating the ten reactions labeled oxidative phosphorylation, with every exchange open, suggested that about 64% of growth depended on the pathway. Taken at face value this was a striking result: it was large and it nominated a candidate anti-angiogenic vulnerability. It also sat awkwardly beside direct functional evidence that complex III is required for endothelial proliferation [20]. Because a clean result that conflicts with independent measurement is a reason for more scrutiny rather than less, we audited the ten reactions against the stoichiometric equations in the reconstruction itself.

Four of the ten reactions (MAR03975, MAR03977, MAR03979 and MAR06912) were named “diphosphate phosphohydrolase” and catalyze inorganic pyrophosphate hydrolysis (H₂O + PPi → H⁺ + 2 Pi) in the endoplasmic reticulum, cytosol, peroxisome and mitochondrion respectively. That activity is unrelated to the electron transport chain and to ATP synthase, and it is carried by a gene set (lhpp, ppa1b, ppa2, prune) encoding no respiratory subunit. They had been misassigned to the OXPHOS subsystem of the reference reconstruction. Two further observations from the audit support the reclassification independently of the reaction names. First, the assignment is internally inconsistent: the reconstruction contains a fifth inorganic pyrophosphatase, MAR07794 in the nucleus, which carries the subsystem label “Miscellaneous”, so four of the five identical reactions are labeled as oxidative phosphorylation and one is not. Second, the four reactions contribute nothing to respiration on the reconstruction’s own stoichiometry: maximizing ATP production on a glucose and oxygen medium in the unmodified reference model gives 31.5 ATP per unit glucose intact and 31.5 with all four blocked, against 2.0 with the corrected set of six reactions blocked, which is exactly the anaerobic yield. The corrected set therefore carries 94% of the network’s maximum ATP generating capacity and the four pyrophosphatases carry none of it. The six retained reactions are complex I (MAR06921), complex III (MAR06918), complex IV (MAR06914), ATP synthase (MAR06916), an FAD-linked dehydrogenase entry into the quinone pool (MAR06911) and an alternative superoxide generating cytochrome *c* oxidase variant (MAR13081) that is constrained to zero flux in the distributed reconstruction. Blocking the four respiratory complexes individually lowers maximum ATP yield to 2.0 (complex IV), 4.0 (ATP synthase), 6.0 (complex III) and 21.5 (complex I, the residual reflecting the FAD-linked bypass). The full audit is given in Table 3.

To establish what the correction does and does not change, we repeated the titration as a factorial over objective, medium, maintenance demand and reaction set (Fig. 4B; Section 2.11). Under the embryonic medium used throughout this study, complete block of the legacy ten reactions and of the corrected six gave identical growth at every maintenance demand and with both objectives: 16% of growth retained with the generic biomass reaction at NGAM 0, 12.5% at NGAM 5 and none at NGAM 20 or above, and 50%, 37% and 0% respectively with MAR00021, which carries no growth-associated ATP. The four misannotated pyrophosphatases therefore play no part in the oxidative phosphorylation result on this medium, and blocking them alone left 95–100% of growth, because the fifth, nuclear pyrophosphatase carries the hydrolysis instead; blocking all five abolished growth in every condition, as expected for an essential biosynthetic reaction. The magnitude of any dependence on oxidative phosphorylation, on the other hand, is set by the objective and the maintenance demand, ranging from half of growth to all of it.

**Fig. 4.**
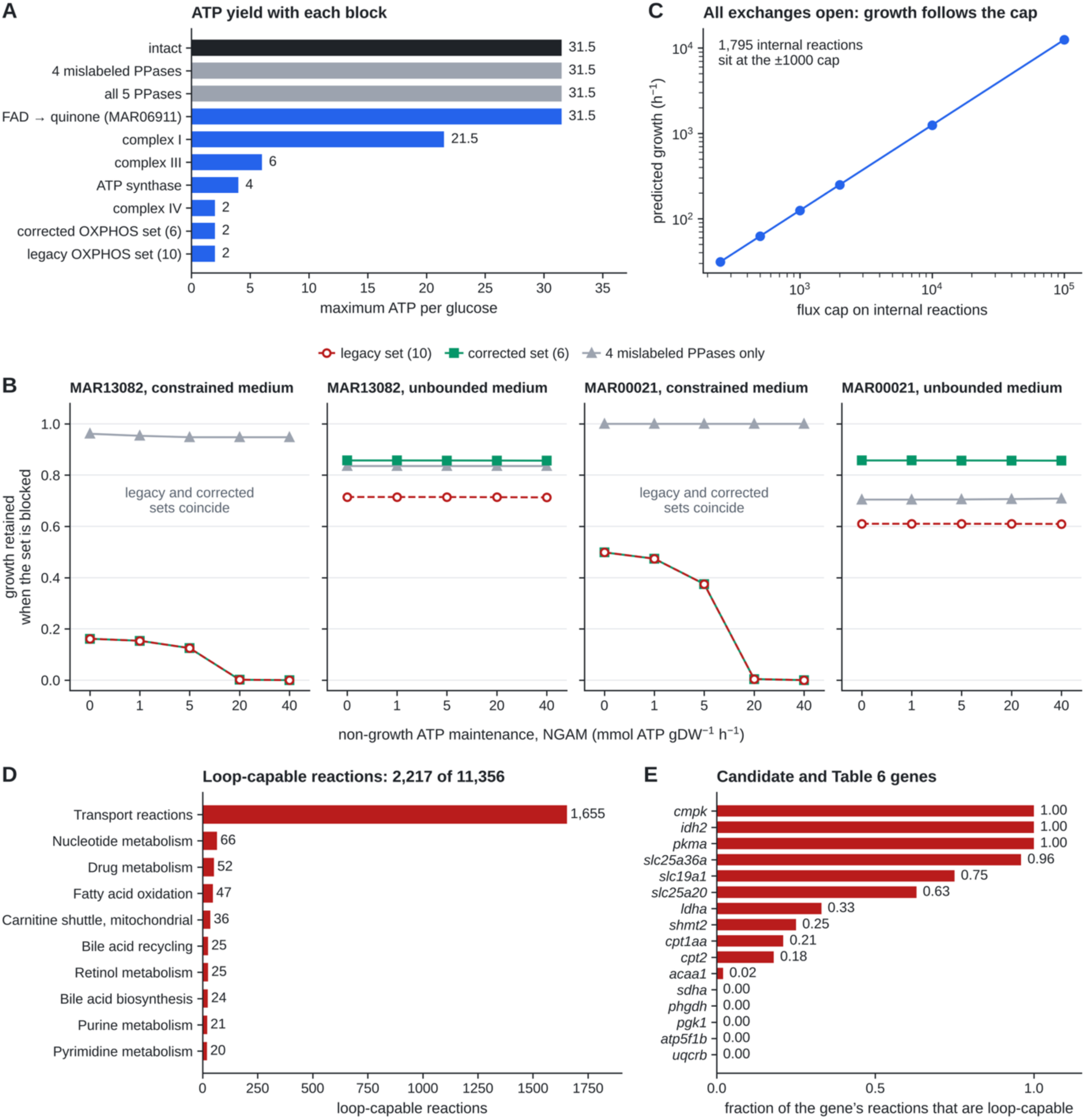
Model audit. **(A)** Maximum ATP yield per glucose in the reference reconstruction, intact and with each reaction set or respiratory complex blocked. **(B)** Growth retained at complete block of the legacy ten-reaction set, the corrected six-reaction set and the four misannotated pyrophosphatases, for two objectives (MAR13082 with 45 ATP growth-associated maintenance; MAR00021 with none), two media (the embryonic medium; all exchanges open) and five non-growth-associated maintenance demands; blocking all five pyrophosphatases gives zero growth in every condition. **(C)** With all exchanges open, growth scales linearly with the bound placed on internal reactions. **(D)** Reactions able to carry flux with every exchange closed, by subsystem. **(E)** Fraction of each gene’s GPR-associated reactions that are loop-capable, for the conventional-screen candidates and the Table 6 targets.

The legacy and corrected definitions diverge only when every exchange is open, where blocking the legacy set retained 61–72% of growth and blocking the corrected set 86%, depending on the objective. That regime is not a usable baseline. With all exchanges open, 1,795 internal reactions sit exactly at their ±1000 bound in the optimum, and growth scales linearly with the bound, from 31 at ±250 to 12,487 at ±100,000 (Fig. 4C); the optimum is therefore set by the arbitrary flux caps rather than by the network or the medium, and the effect of removing the four mislabeled pyrophosphatases there reflects the capped capacity of the remaining nuclear route. We therefore report the annotation error as the robust result of this section, and do not report a quantitative dispensability figure from it. Under the embryonic medium, complete block of the corrected electron transport chain and ATP synthase costs most or all of growth, which is what a network with no alternative high-yield ATP source should show, and is consistent with the functional requirement for complex III in proliferating endothelium [20].

The general lesson is that a pathway level dispensability result is a statement about a reaction set, a medium, an objective and a maintenance demand jointly, and is uninterpretable if any of them is left implicit; a result obtained with every exchange open should in addition be checked for dependence on the internal flux bounds.

### 3.5. One fifth of the reference network can carry flux in internal cycles, including every candidate of the conventional screen

With every exchange, biomass and maintenance reaction closed, 2,217 of the 11,356 flux-consistent reactions (19.5%) could still carry flux, and are therefore members of thermodynamically infeasible cycles (Fig. 4D). Most are transport reactions (1,655), with smaller groups in nucleotide metabolism, fatty acid oxidation and the mitochondrial, cytosolic and endoplasmic carnitine shuttles. The candidates of the conventional screen are among the most exposed (Fig. 4E): 64 of the 102 reactions associated with slc25a20, 22 of 23 for slc25a36a, the single reaction of idh2, 12 of 16 for slc19a1 and all 41 for cmpk, together with all seven pyruvate kinase reactions and both lactate dehydrogenase reactions. Loops do not affect the pFBA solutions used here, which were within 10⁻⁷ of their nearest loop-free flux in every model, because minimizing total flux removes them. They do affect flux variability ranges and any procedure that pushes fluxes to vertices of the feasible space: the range of the carnitine– acylcarnitine carrier at 90% of optimum fell from −18.8 to 2.5 to −2.5 to 2.5 once loops were removed, and the lactate dehydrogenase, CPT1A and GLUT1 ranges were also loop-dependent (Table 6). With bounds scaled from ±1000, as in the conventional screen, such reactions reach ranges of exactly ±50 (the floor of 0.05 times 1000), the signature of flux running round a loop to an expression bound.

### 3.6. Expression constraints that bind, and flux level tests on metacell models

Expression-scaled bounds constrain a network only if they are comparable to the fluxes the medium allows. When bounds were scaled from the default of ±1000, as is common, the median state model had 8 reactions at an expression bound and retained 87% of the growth of the unconstrained network; with bounds scaled from a capacity of 50, the median model retained 47% of growth and 33 reactions were bound (Fig. 5A). We used U = 50 as the primary setting and U = 100 as a sensitivity setting, and built a separate model for each of 325 metacells of about 20 cells across the endothelial states and five non-endothelial lineages.

**Fig. 5.**
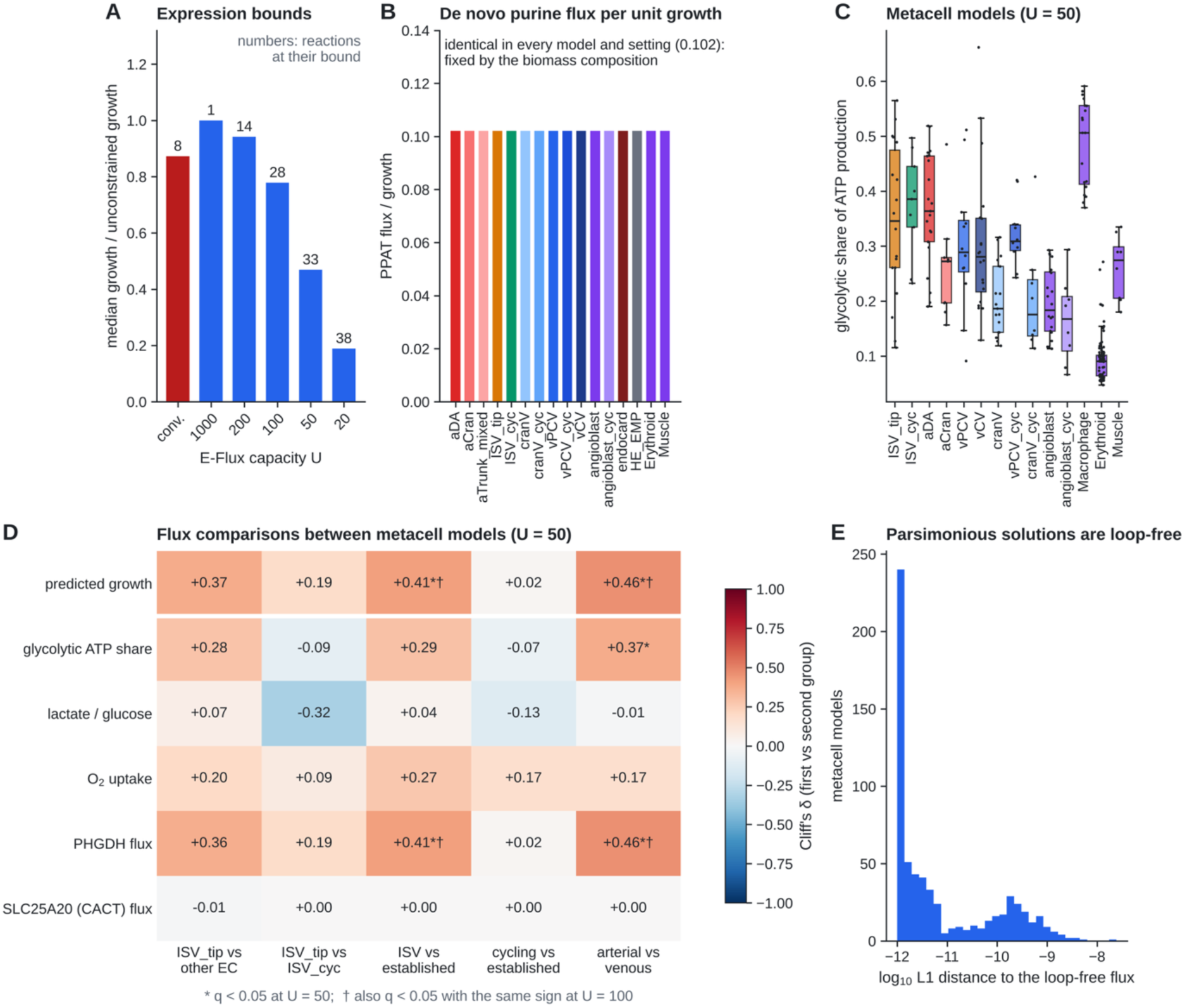
Expression constraints and flux level tests. **(A)** Median growth relative to the unconstrained network, and median number of reactions at their expression bound (bar labels), for expression-scaled bounds with capacities from 1000 to 20 and for the conventional scaling (normalization to the maximum, U = 1000; “conv”). **(B)** De novo purine synthesis flux per unit growth in each state model (U = 50). **(C)** Glycolytic share of ATP production in metacell models (U = 50). **(D)** Flux level comparisons between compartments, as Cliff’s delta, with * marking q < 0.05 at U = 50 and † significance with the same sign at U = 100 (Benjamini–Hochberg across all flux level tests). **(E)** L1 distance between each pFBA solution and its nearest loop-free flux distribution. The first row is predicted growth; PHGDH flux is a fixed multiple of it.

Two read-outs behaved as the structure of the model dictates rather than as biology might suggest. De novo purine synthesis per unit growth was 0.102 in every model under every setting (Fig. 5B), because in a growth-maximizing model biosynthetic fluxes are fixed by the composition of the biomass reaction, so they can rank cell states only by predicted growth. A flux level test of nucleotide or one-carbon claims of the kind applied in the conventional screen therefore cannot fail or succeed on the biology, and the marker level results of Section 3.3 are the only evidence these data offer on them. The glycolytic share of ATP production, by contrast, varied between states (Fig. 5C), and it ranked lineages plausibly, macrophages highest (median 0.51) and erythroid cells lowest (0.09).

Among endothelial compartments, the glycolytic share of ATP was higher in sprouting ISV models than in established-vessel models (median 0.36 against 0.28, Cliff’s δ = 0.29) but not significantly so after correction (q = 0.11), and did not differ between ISV_tip and ISV_cyc (δ = −0.09, q = 0.96; Fig. 5D). The one difference between sprouting and established vessels significant at both capacities was predicted growth (δ = 0.41, *q* = 0.02 at U = 50; δ = 0.37, *q* = 0.04 at U = 100). Phosphoglycerate dehydrogenase flux showed the identical difference, because in every sprouting and established-vessel metacell it was a fixed multiple of growth (0.0012), as purine synthesis was (Fig. 5D). The models therefore predict a higher biosynthetic capacity in sprouting vessels, which serine synthesis follows, rather than a separate routing of carbon into serine; the serine and one-carbon marker score was not higher in sprouting cells (Table 5). Predicted growth, and with it PHGDH flux, was also higher in arterial than in venous models at both capacities, and the glycolytic share was higher in arterial models at U = 50 only. At the flux level, then, the tip glycolysis claim is not supported as a tip-versus-stalk difference, and the sprouting-versus-established difference it reflects is a trend.

### 3.7. Candidate vulnerabilities do not survive an ensemble of extraction methods and a full lineage control

A gene whose deletion impairs sprouting endothelium is a useful candidate only if the call does not depend on how the network was extracted and if it spares other tissues. We therefore deleted each of the 825 genes whose loss disables at least one reaction in every model of a four-member ensemble (FASTCORE at three thresholds, and GIMME), across all fourteen endothelial and hemogenic states and all thirteen non-endothelial lineages (Fig. 6A). Under the two-lineage rule, which requires inhibition of sprouting growth by at least 0.10 while erythroid and muscle models are spared, the four members called 13, 11, 5 and 5 genes, and the lists barely overlapped: the loose and default FASTCORE members called the succinate dehydrogenase subunits and other TCA cycle genes (sdha–d, suclg1, dlst, fh, dldh), the strict member a different set (acat2, cpt2, lcat, nmnat1, slc25a19), and GIMME a third (fdps, idh2, mdh2, ptdss1a, slc25a36a). Under the all-lineage rule, which in addition requires sparing of all thirteen non-endothelial lineages and a within-endothelium selectivity of at least 0.10, the FASTCORE loose and default members called no gene, the strict member called lcat and slc25a19, and GIMME called slc25a36a (Fig. 6B, Table 4). No gene was called by more than one member. The carnitine–acylcarnitine carrier slc25a20, the lead candidate of the conventional screen, had no effect on growth in any FASTCORE model at any threshold and inhibited sprouting growth by 0.05 under GIMME.

**Fig. 6.**
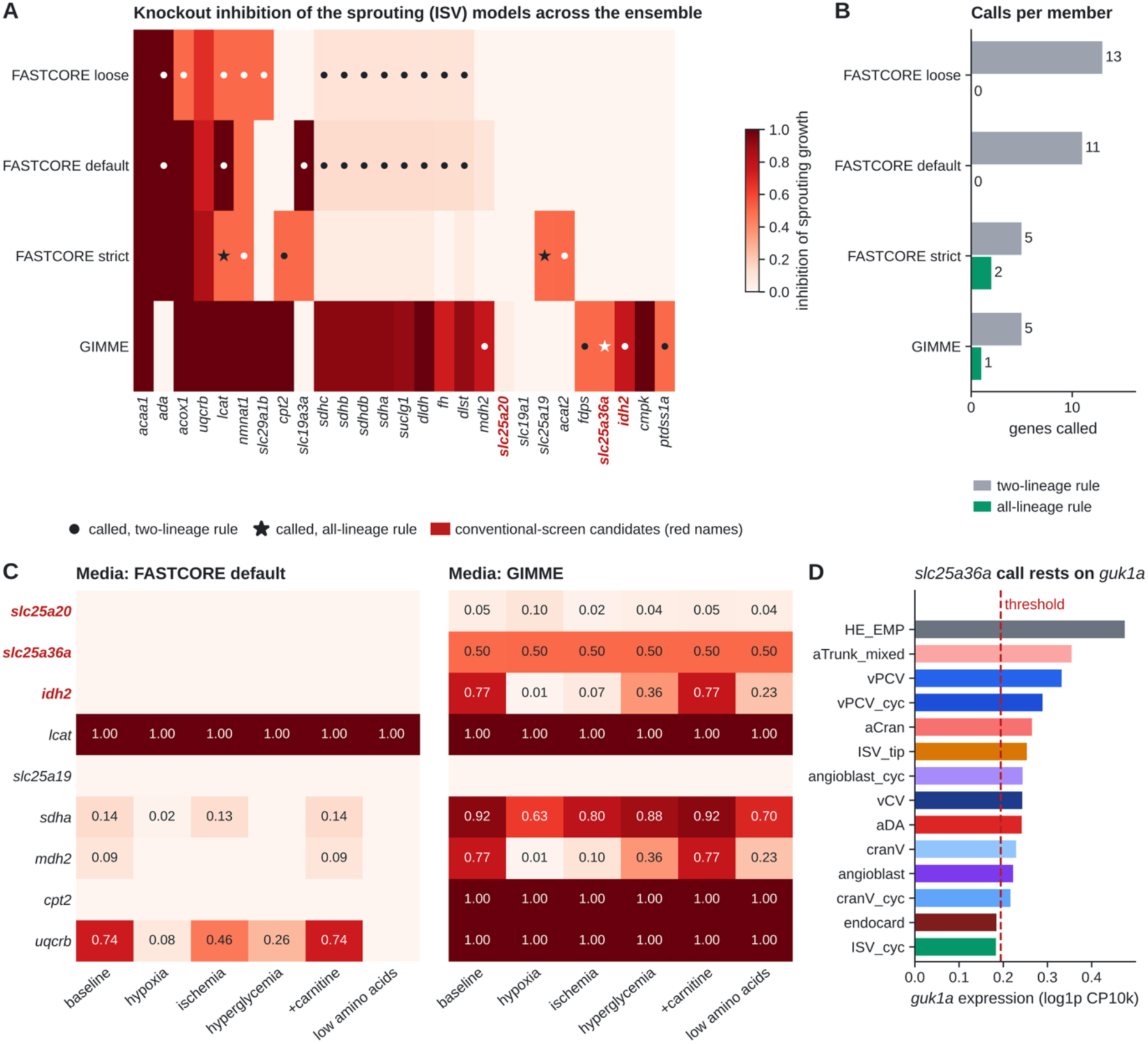
Knockout screen across a four-member model ensemble. **(A)** Inhibition of growth in the sprouting models (mean of ISV_tip and ISV_cyc) after deletion of each gene called in any member or nominated by the conventional screen (red labels), for FASTCORE at three local-T2 thresholds and GIMME; • called under the two-lineage rule (sparing of erythroid and muscle models), ★ called under the all-lineage rule (sparing of all thirteen non-endothelial lineages and within-endothelium selectivity of at least 0.10). **(B)** Number of genes called per member under each rule. **(C)** Sprouting inhibition across media for the FASTCORE default and GIMME members. **(D)** Expression of guk1a across endothelial states against the GIMME threshold (dashed); its omission from the ISV_cyc model is the sole cause of the slc25a36a call.

**Table 4.**
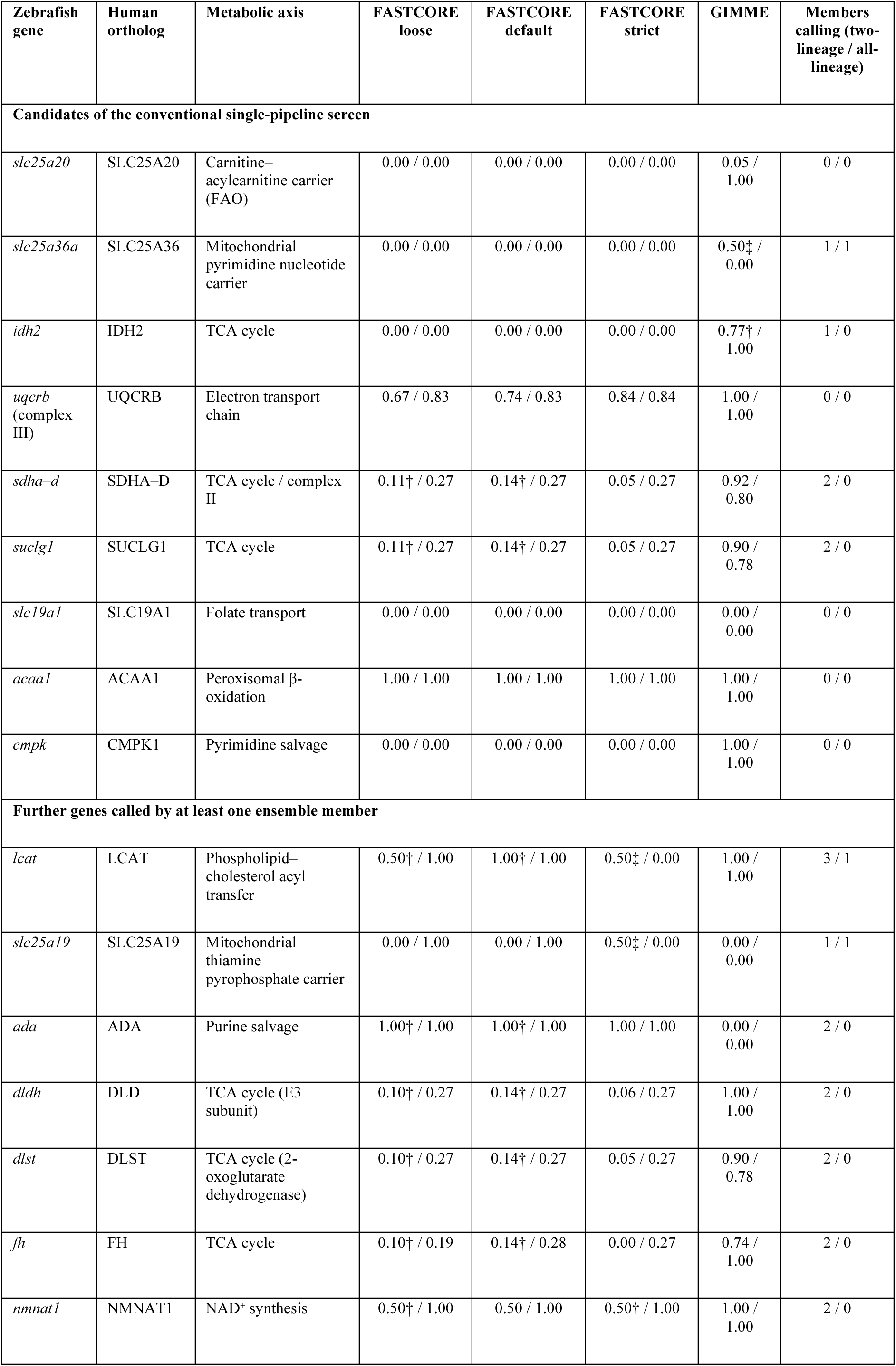
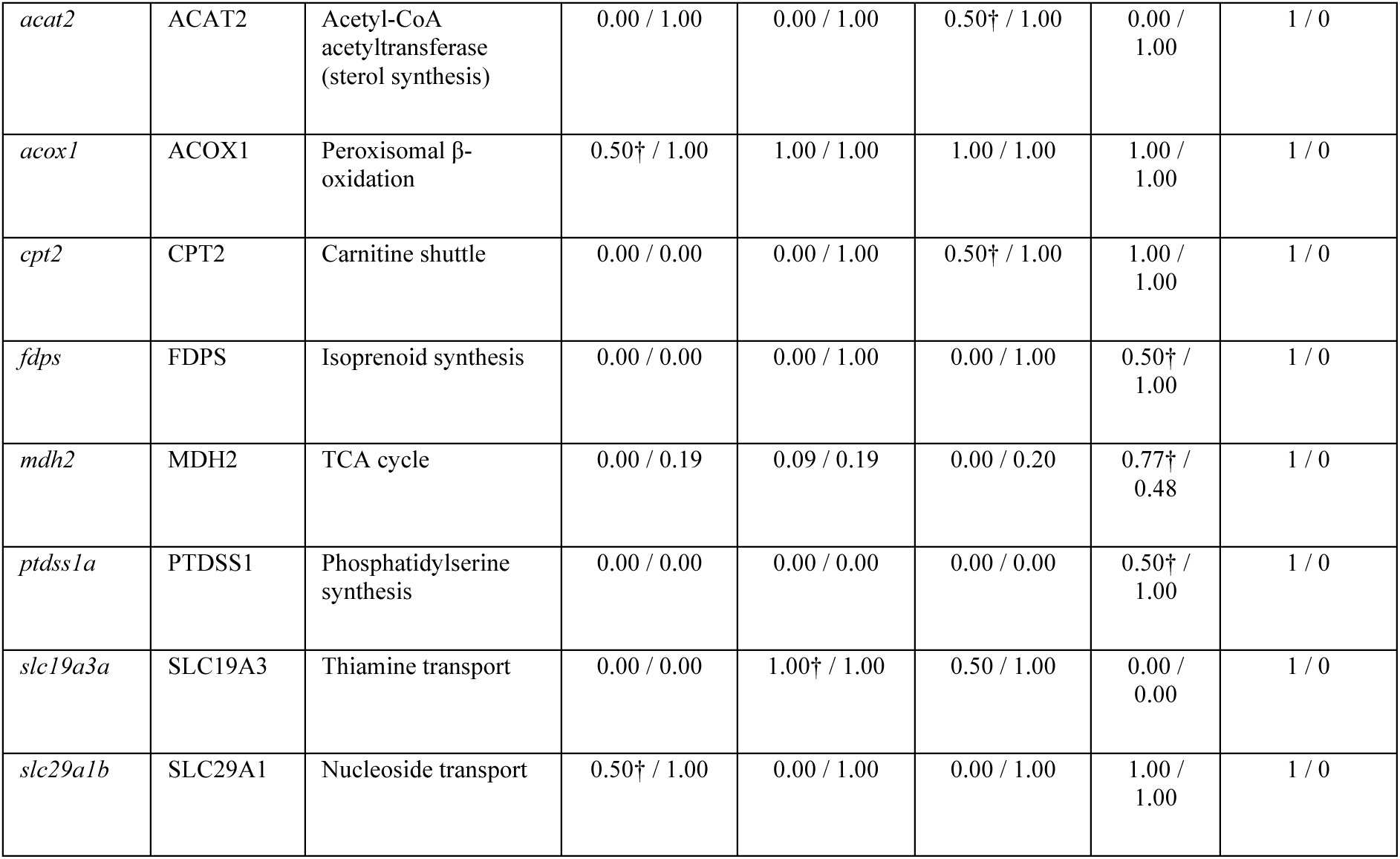
Robustness of candidate calls across a four-member model ensemble. Each cell gives the predicted inhibition of sprouting endothelium (mean of ISV_tip and ISV_cyc; 0, no effect; 1, lethal), followed by the largest inhibition in any of the 13 non-endothelial lineages, for FASTCORE models built at three local-T2 threshold settings (loose, 15th/65th; default, 25th/75th; strict, 40th/85th percentiles) and for GIMME. A value of 0.50 means the knockout is lethal in one of the two sprouting models. † Called under the two-lineage rule: sprouting inhibition of at least 0.10 and no inhibition in the erythroid or skeletal muscle model. ‡ Called under the all-lineage rule: sprouting inhibition of at least 0.10, no inhibition in any non-endothelial lineage, and sprouting minus established-vessel inhibition of at least 0.10. The upper block lists the candidates of the conventional single-pipeline screen (Supplementary Note S1); the lower block lists every further gene called by at least one member. SDHA–D share one row because their knockouts are identical. Each all-lineage call traces to a single pruning difference between the two sprouting models: *slc25a36a* (GIMME) is lethal in ISV_cyc because cytosolic guanylate kinase (MAR04020) was omitted when *guk1a* fell 6% below its threshold (0.183 against 0.194 log1p CP10k); *slc25a19* (FASTCORE strict) is lethal in ISV_tip because uridine phosphorylase (*upp1*/*upp2*) was pruned from that model only; *lcat* (FASTCORE strict) depends on an 81-reaction endoplasmic reticulum carnitine shuttle route retained in ISV_tip and not in ISV_cyc. Values are model predictions.

**Table 5.** Metabolic claims tested at the marker level and the flux level. Marker level: difference in per-cell pathway score between the focal compartment and the comparison, with one-sided Mann–Whitney q values (Benjamini– Hochberg across claims), raw and adjusted for S and G2/M scores and sequencing depth (ordinary least squares with HC3 errors). Flux level: Cliff’s δ between metacell models (about 20 cells each; 325 models) at two E-Flux capacities, U = 50 and U = 100, with Benjamini–Hochberg q values. A claim is supported at the flux level only when q < 0.05 in the stated direction at both capacities. Compartments: tip, ISV_tip; cycling, ISV_cyc and the cycling venous, cranial venous and angioblast states; arterial, aDA and aCran; venous, vPCV, vCV and cranV; established vessels, arterial plus venous. ^a^ Comparison outside the eight pre-specified claims; nominal p values, not included in the Benjamini–Hochberg set. ^b^ In every ISV and established-vessel metacell, de novo purine flux (PPAT) and serine synthesis flux (PHGDH) are fixed multiples of predicted growth (0.102 and 0.0012), so differences in these fluxes reflect differences in predicted growth; the PHGDH δ equals the growth δ at both capacities. All p values are computed on cells from a single pooled clutch and quantify within-sample separation only.

| Claim | Source | Comparison | Marker $\Delta$, raw (q) | Marker $\Delta$, adjusted (q) | Flux readout | Flux $\delta$, $U = 50$ (q) | Flux $\delta$, $U = 100$ (q) | Verdict |
| --- | --- | --- | --- | --- | --- | --- | --- | --- |
| Tip cells are the most glycolytic endothelial cells | [6] | Tip vs other endothelium | +0.034 ($5 \times 10^{-10}$) | +0.037 ($5 \times 10^{-11}$) | Glycolytic ATP fraction | +0.28 (0.20) | +0.27 (0.22) | Marker level only |
| Tip cells are more glycolytic than stalk cells | [6] | Tip vs cycling ISV | +0.005 (0.61) | +0.007 (0.69) | Glycolytic ATP fraction | −0.09 (0.96) | +0.02 (1.00) | Not supported |
| Sprouting vessels are more glycolytic than established vessels | This study | ISV (tip + cycling) vs established | +0.027 ($p = 6 \times 10^{-7}$)<sup>a</sup> | +0.027 ($p = 2 \times 10^{-7}$)<sup>a</sup> | Glycolytic ATP fraction | +0.29 (0.11) | +0.16 (0.50) | Marker level only |
| The arterial lineage is the most glycolytic | Lineage-level | Arterial vs other endothelium | +0.028 ($3 \times 10^{-9}$); | +0.032 ($2 \times 10^{-12}$); | Glycolytic ATP fraction | +0.37 (0.04) | +0.19 (0.45) | Not supported |
|  | restatement of [6] |  | tip scores highest | tip scores highest | (arterial vs venous) |  |  |  |
| Proliferating endothelium synthesizes nucleotides | [10] | Cycling vs other endothelium | +0.022 ($7 \times 10^{-31}$) | +0.018 ($1 \times 10^{-17}$) | PPAT flux per unit growth (cycling vs established) | +0.12 (0.58) | -0.16 (0.48) | Marker level only; flux fixed by biomass composition<sup>b</sup> |
| Proliferating endothelium uses serine / one-carbon metabolism | [13] | Cycling vs other endothelium | +0.011 ($5 \times 10^{-6}$) | +0.013 ($6 \times 10^{-5}$) | PHGDH flux (cycling vs established) | +0.02 (1.00) | +0.03 (0.96) | Marker level only |
| Serine synthesis is higher in sprouting vessels | This study | ISV (tip + cycling) vs established | -0.004 (p = 0.88)<sup>a</sup> | -0.005 (p = 0.05)<sup>a</sup> | PHGDH flux | +0.41 (0.023) | +0.37 (0.038) | Flux level only; follows predicted growth<sup>b</sup> |
| Quiescent endothelium upregulates fatty acid β-oxidation | [11] | Venous vs other endothelium | +0.010 ($3 \times 10^{-4}$) | +0.003 (0.18) | CACT flux | Zero in all models | Zero in all models | Not supported after adjustment |
| TCA cycle activity is highest in cycling endothelium | Conventional screen | Cycling vs other endothelium | -0.006 (1.00) | -0.007 (1.00) | — | — | — | Not supported |
| Tip cells have the highest OXPHOS activity (corrected gene panel) | Conventional screen | Tip vs other endothelium | +0.056 ($5 \times 10^{-15}$) | +0.058 ($4 \times 10^{-17}$) | O<sub>2</sub> uptake | +0.20 (0.45) | +0.09 (0.58) | Marker level only |
| The carnitine shuttle is elevated in cycling endothelium (SLC25A20 axis) | Conventional screen | Cycling vs other endothelium | -0.028 (1.00) | -0.015 (1.00) | CACT flux | Zero in all models | Zero in all models | Opposite direction |

Tracing the calls to their cause showed why they were not robust. Expression of slc25a20, slc25a36a and idh2 offers no basis for selectivity: slc25a20 is detected at similar levels in every lineage, and idh2 is higher in neural progenitors, fibroblasts, mesenchyme and muscle than in any endothelial state. The GIMME call of slc25a36a rests on a single reaction. The knockout is lethal in ISV_cyc and tolerated in ISV_tip, and the smallest set of reactions from the ISV_tip model that restores growth in ISV_cyc is one cytosolic guanylate kinase reaction (MAR04020). GIMME omitted it from ISV_cyc because guk1a expression there (0.183, log-normalized) falls just below the extraction threshold (0.194), while it lies above in ISV_tip (0.254) and in most other endothelial states (Fig. 6D). The strict-threshold FASTCORE call of slc25a19 is of the same kind, lethal in ISV_tip only because uridine phosphorylase (upp1 or upp2) was pruned from that model, and the lcat call depends on an endoplasmic carnitine-shuttle route retained in one sprouting model and not the other. Each of these is a statement about which reactions an extraction retained, not about the metabolism of the cell state.

Because the metabolic environment of a sprouting vessel differs between developmental and disease settings, we repeated the screen for the union of called genes under three disease media, with carnitine supplementation and with a tenfold lower amino acid supply (Fig. 6C). These media differ from baseline only in the named exchange bounds and are a sensitivity analysis, not a model of a disease microenvironment. Calls that were stable across media were stable because they were structural (lcat, cpt2, slc25a36a under GIMME), while others changed with the medium: idh2 inhibition under GIMME fell from 0.77 at baseline to 0.01 under hypoxia, and complex III (uqcrb) inhibition in the default FASTCORE member ranged from 0.74 to 0(low amino acid supply). Carnitine supplementation changed no call.

### 3.8. Recapitulation of published endothelial metabolism phenotypes

As an external check that is independent of the claims tested in Section 3.3, we compiled eighteen endothelial metabolism phenotypes from the experimental literature and asked whether the models reproduce each one under a rule stated in advance (Section 2.10): a phenotype counts as recapitulated if predicted flux through the target is higher in the sprouting models than in the established-vessel models, or if deleting the target reduces sprouting growth to 0.90 or below (Table 6). Under that rule, eleven of the eighteen were recapitulated, all of them by the flux criterion: the carnitine–acylcarnitine carrier SLC25A20, succinate dehydrogenase, CPT1A, acetyl-CoA carboxylase, fatty acid synthase, pyruvate kinase, ATP synthase, ribonucleotide reductase, SHMT2, hexokinase and PHGDH. The criterion is, however, weaker than it appears. The pooled sprouting model grows 1.26 times faster than the pooled established-vessel model, and five of the eleven flux ratios (CPT1A, fatty acid synthase, ribonucleotide reductase, SHMT2 and PHGDH) equal that growth ratio exactly, because their flux is drawn by biomass synthesis; ATP synthase, at 1.14, carries proportionally less flux in the sprouting model than growth alone would predict. After division by growth, flux was elevated in the sprouting model for only five targets: succinate dehydrogenase, acetyl-CoA carboxylase, pyruvate kinase, hexokinase and SLC25A20. The last does so on a loop-dependent range, which narrowed from −18.8 to 2.5 to −2.5 to 2.5 once loops were removed, and its deletion had no effect on growth in any of the four ensemble members.

**Table 6.**
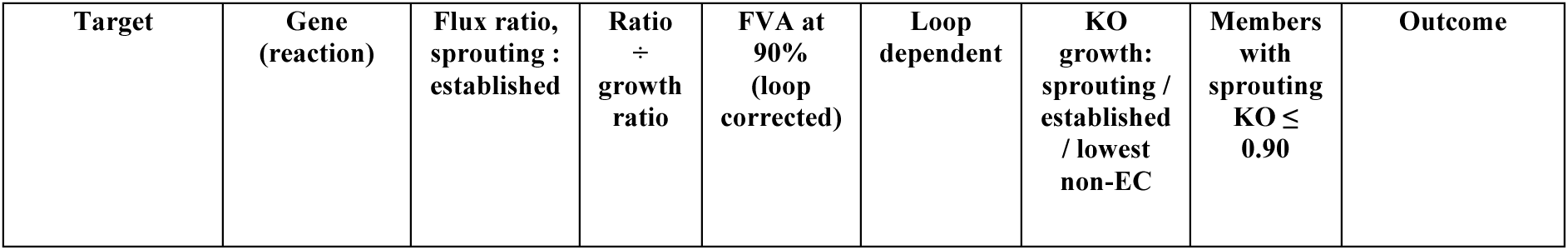
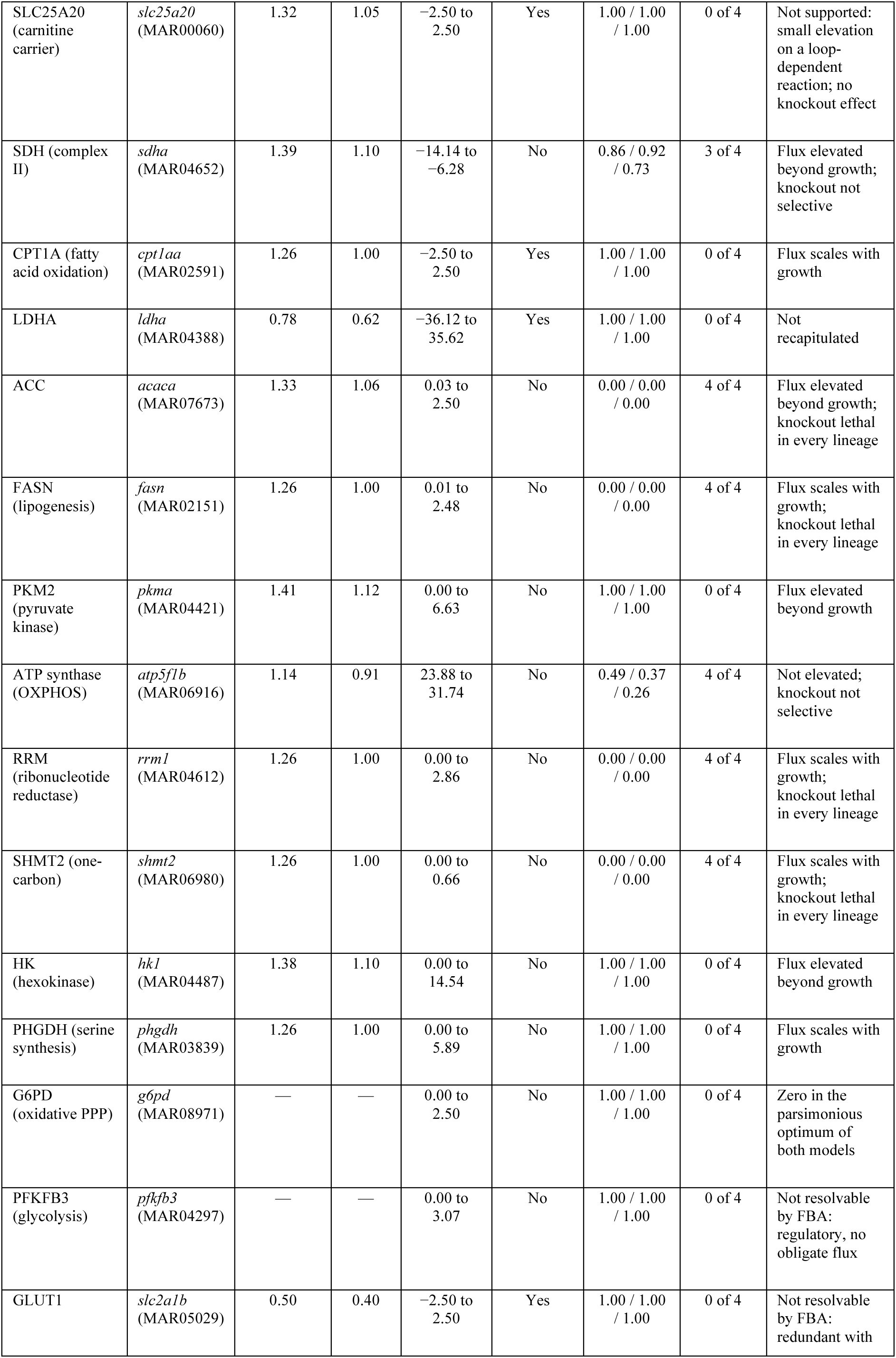
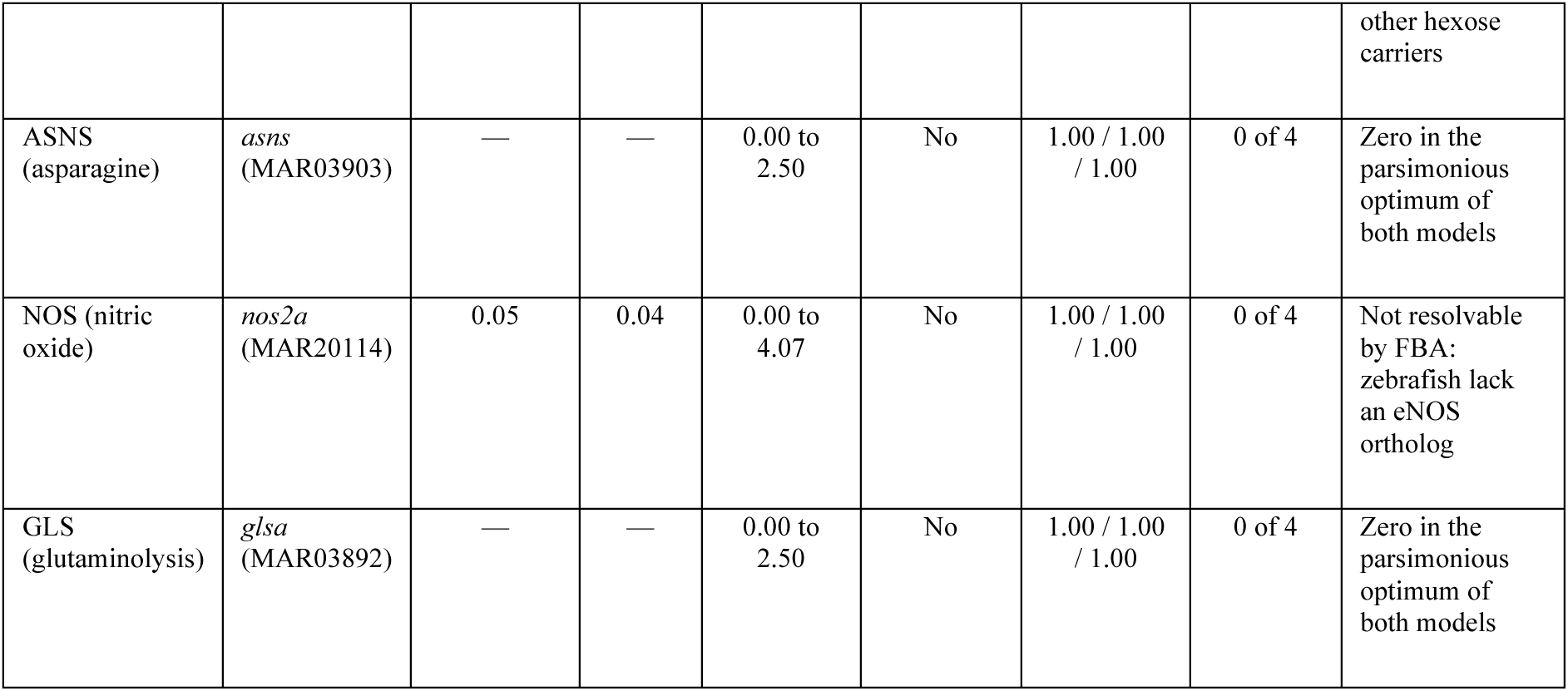
Published endothelial metabolism phenotypes recomputed with loop correction. Sprouting (ISV_tip and ISV_cyc) and established-vessel (aDA, aCran, vPCV, vCV and cranV) pseudobulk E-Flux models (U = 50). Flux ratio: summed absolute flux over the gene’s reactions in the loop-corrected parsimonious solution, sprouting over established; a dash marks zero flux in both models. Predicted growth is 1.26 times higher in the sprouting model, so a ratio of 1.26 (1.00 after division) means that the flux only follows growth. FVA at 90% of optimum on the gene’s highest-flux reaction, after loop correction; “loop dependent” marks reactions whose range shrinks when thermodynamically infeasible cycles are removed. Knockout (KO) growth is relative to the unperturbed model in the FASTCORE default member, for the sprouting models, the established-vessel models and the most affected of the 13 non-endothelial lineages; the next column counts ensemble members (of four) in which the sprouting knockout leaves at most 0.90 of growth. Every target is present in Zebrafish-GEM; NOS is represented by *nos2a* because zebrafish lack an eNOS ortholog. Four targets show flux elevated beyond growth (SDH, ACC, PKM2 and HK); six reduce sprouting growth in at least three members, and none of these spares the non-endothelial lineages.

The deletion criterion separates targets more sharply. Six targets reduced sprouting growth to 0.90 or below in at least three of the four ensemble members, but none did so selectively. Four of them (acetyl-CoA carboxylase, fatty acid synthase, ribonucleotide reductase and SHMT2) are lethal in every lineage we modeled, and are general biosynthetic requirements rather than endothelial phenotypes. ATP synthase inhibits non-endothelial models more than sprouting ones, the lowest non-endothelial growth after deletion being 0.26 against 0.49 in sprouting endothelium. Succinate dehydrogenase inhibits sprouting endothelium modestly (0.86 of growth retained) but some non-endothelial lineages more (0.73). No target therefore reproduces its published phenotype with selectivity for sprouting endothelium, and we regard this, rather than the count of eleven, as the honest summary of how far these models reproduce what has been measured.

The seven phenotypes that were not recapitulated fall into three interpretable classes, and we report them in full because their failure modes are informative about what this class of model can and cannot do. LDHA carries less flux in the sprouting model than in the established-vessel model (ratio 0.78), on a feasible range that is itself loop-dependent. Three (G6PD, asparagine synthetase and glutaminase) carry no flux in either parsimonious solution, yet each has a non-zero feasible range at 90% of optimum, so the phenotype is reachable in the network but is not selected under minimum-flux parsimony; for glutaminase this also reflects the supply of glutamate from other routes and from the medium. Three (PFKFB3, GLUT1 and nitric oxide synthase) are not resolvable by growth-coupled flux balance analysis at all. The critical case is PFKFB3: it is the founding target of the field, and it cannot be recovered by this method because it is a regulatory enzyme that sets the glycolytic rate through fructose-2,6-bisphosphate rather than carrying obligate pathway flux itself. The same argument applies to GLUT1, whose transport function is redundantly covered by other hexose carriers, and to nitric oxide synthase, which zebrafish lack in its endothelial isoform and whose product the biomass reaction does not require. These are limitations of constraint-based modeling as a formalism rather than defects of this particular reconstruction, and a validation exercise that reported only the successes would misrepresent what the method can do.

## 4. Discussion

### 4.1. A flux-resolved view of endothelial metabolic heterogeneity

This study set out to test whether the foundational tip, stalk and venous metabolic hypotheses of endothelial biology hold when re-examined at single-cell resolution with formal statistical validation and network-level flux constraints. The answer is mixed, and in one respect different from the question as posed. In these data the tip-like and stalk-like populations of the intersegmental vessel are one sprouting program sampled in S and in G2/M, so the tip/stalk contrast could not be tested as such; what could be tested is the contrast between sprouting and established vessels. Glycolytic elevation, the founding claim of the field, held as a difference between the sprouting intersegmental vessel and established arterial and venous endothelium at the marker level, with and without control for cell-cycle phase, but was only a trend at the flux level and was absent within the sprouting population itself. Elevation of nucleotide and serine/one-carbon gene expression in cycling endothelium was supported at the marker level, whereas venous fatty acid oxidation was not once cell-cycle phase and sequencing depth were controlled. The one flux-level difference that held at both capacities tested was a higher predicted growth capacity in sprouting than in established vessels, with serine synthesis through phosphoglycerate dehydrogenase rising in fixed proportion; this is compatible with, but does not by itself demonstrate, the requirement for PHGDH in vessel formation [13].

We read these results not as a refutation of the underlying experiments, which measured quantities our models do not, but as a statement about what a constraint-based model can and cannot corroborate. In a growth-maximizing model the flux through a biosynthetic pathway is fixed by the composition of the biomass reaction, so nucleotide and one-carbon synthesis can differ between models only through predicted growth, and a flux-level test of claims about them can neither fail nor succeed on the biology; the marker level is the only evidence these data offer on them. The practical consequence for anyone building on this literature is that the statistical strength of a marker-level result is a poor guide to whether it will behave as a flux phenotype, and that the two levels must be tested separately, with the structure of the objective in mind, and neither taken as evidence for the other. The same caution applies to the distinction between lineage and state: the claim that arterial cells are the most glycolytic population failed at the whole-lineage level while the sprouting vessel scored highest, which is exactly the inversion that averaging over a heterogeneous compartment produces and that single-cell resolution exists to prevent. Single-cell resolution brings its own confound, however. Proliferation separates cells as strongly as identity does, and a state boundary drawn by clustering should be checked against cell-cycle phase before it is read as a division of labor.

### 4.2. Reaction-annotation risk as a methodological hazard

A consequential methodological result of this work concerns the reaction lists that pathway analyses rely on. Four of the ten reactions labeled oxidative phosphorylation in Zebrafish-GEM are inorganic pyrophosphatases, which is evident from their stoichiometry and gene associations alone and inconsistent with a fifth, identical reaction labeled Miscellaneous. In the conventional screen the subsystem label produced an apparent 64% growth dependence on the pathway, and the factorial of Section 3.4 shows what that figure measured. It arose only with every exchange open, a regime in which growth is set by the arbitrary bounds on internal reactions, and in which removing four of the five pyrophosphatases limits the capacity of the remaining route for an essential biosynthetic reaction. Under a defined medium the misannotation plays no part, and dependence on the corrected pathway is set by the objective and the maintenance demand, between half of growth and all of it. A misassigned reaction is not merely noise in a pathway score; combined with an unconstrained medium, it can make a coherent, mechanistically plausible and entirely spurious dependence.

This risk is structural rather than incidental. Zebrafish-GEM is a derivative of Human-GEM obtained by orthology mapping, so a single subsystem assignment error propagates into every derived model and every species that inherits it, and it is invisible to any workflow that selects reactions by subsystem label, which is the standard practice, including in comparable network analyses across human tumors. The error is also of a particularly difficult kind to detect: the subsystem did contain genuine electron transport reactions, so a superficial check found real biology, and the spurious result agreed with a plausible prior, so it felt corroborated rather than suspicious. That the same reconstruction labels a fifth, identical pyrophosphatase “Miscellaneous” shows the inconsistency is detectable from the model file alone, by anyone who looks. We suggest four checks as a standard step before a constraint-based study reports a selective vulnerability claim, and we have applied all four here: reaction by reaction inspection of stoichiometry and gene association against subsystem labels; an audit of thermodynamically infeasible cycles, with flux ranges and ratios reported for loop-capable reactions only after loop correction; a test at two levels in which a claim must survive both marker statistics, with cell-cycle phase controlled, and flux level comparison across replicate models; and a specificity control in which every candidate is evaluated across an ensemble of extraction methods and thresholds and in every non-target lineage built by the same pipeline. To these we would add an explicit statement of the objective, the medium and the nutrient that limits growth on it, and the maintenance demand, together with a check that expression-scaled bounds actually bind, since Sections 3.4 and 3.6 show that each of these can govern a result as much as the reaction list does. These complement rather than replace existing model quality and reproducibility standards.

### 4.3. Why vulnerability calls from a single extraction fail, and what remains testable

The conventional single-pipeline screen (Supplementary Note S1) nominated the carnitine– acylcarnitine carrier SLC25A20, the mitochondrial pyrimidine nucleotide carrier SLC25A36 and IDH2 as selective vulnerabilities of sprouting endothelium. None survived the ensemble, and the reason is instructive and, we suspect, general. None of the three genes is preferentially expressed in endothelium, so any selectivity must come from which alternative reactions an extraction retains in each model, and which reactions are retained depends on where each gene sits against a threshold. The one call that met the all-lineage rule under GIMME is a clean example: the slc25a36a knockout is lethal in the cycling intersegmental vessel model only because a single guanylate kinase reaction was omitted from it, and it was omitted because guk1a expression in those cells fell 6% below the extraction threshold. A within-endothelium ranking computed on that one extraction would have reported a sprouting-selective vulnerability, and a control against two lineages would not have caught it. We therefore propose that a candidate from a context-specific screen be reported only if it is called across an ensemble of extraction methods and thresholds, and only with the reactions that make it lethal identified.

These results do not show that the mitochondrial carriers are irrelevant to angiogenesis; they show that these models give no evidence that they are. The question is open to a direct test. A homozygous slc25a20 zebrafish mutant has been described in which most homozygotes survive embryogenesis but die before adulthood, and no vascular phenotype has been reported [65]. Imaging intersegmental vessel sprouting in that line, with attention to maternally supplied transcript, would test the conventional screen’s prediction at little cost. SLC25A20 deficiency in humans causes a severe metabolic disorder with cardiac and hepatic involvement [66], so carnitine-shuttle inhibition should not in any case be assumed to be endothelium-sparing. Metabolic rather than VEGF-directed targeting remains attractive given the resistance and toxicity that anti-VEGF therapy encounters [19], and the higher biosynthetic capacity predicted for sprouting vessels is compatible with the experimentally established PHGDH dependence of endothelium [13], but the present models do not nominate a selective target, and we do not claim one.

### 4.4. Limitations and future directions

Several limitations warrant explicit acknowledgment. First, the data derive from a single pooled clutch sorted into two libraries at one developmental stage. Every per-cell test therefore establishes that a metabolic difference is real within this sample and not that it would reproduce in an independently raised clutch, and the metacell models, which partition the same cells, cannot recover between-sample variance; replication in an independent zebrafish endothelial dataset, such as the Daniocell [67] or Zebrahub [68] atlases, is the most direct remedy. Tip and stalk are here transcriptional states rather than live migratory behaviors, and a developmental time course with independent biological replicates is a natural extension. Second, in these data the two intersegmental vessel populations differ mainly in cell-cycle phase, so tip-versus-stalk claims could not be tested as such; a dataset with spatial or lineage information, or one from a later stage at which stalk cells are more numerous, is needed for that. Third, E-Flux constrains reaction bounds by expression without pruning the network, its effect depends on a capacity that has to be calibrated (Fig. 5A), and although the parallel context-specific models address this, neither captures enzyme kinetics, allostery or post-translational regulation, which is why a regulatory enzyme such as PFKFB3 is not resolved by growth-based essentiality (Section 3.8), and why coupling constraint-based models to structural and kinetic simulation is a natural next step. Fourth, the oxidative phosphorylation factorial was run on the reference network, and the state models use a single objective and maintenance demand (NGAM = 1); Section 3.4 shows that the size of any dependence on energy pathways moves with both. Growth on the embryonic medium is set by amino acid supply, so results concerning carbon and oxygen supply should be read with that in mind.

Fifth, no gene met the all-lineage rule in more than one ensemble member, so this study reports no selective candidate; the pathway level findings, and the checks themselves, are its primary output. Sixth, about half of an extracted network depends on the algorithm (mean Jaccard index 0.48 between FASTCORE and GIMME models of the same group); the ensemble spans two algorithms and three thresholds, and other extraction methods, such as INIT [38] or iMAT [69], could change individual calls. Seventh, the objective is growth maximization throughout, including for the tip state, which is canonically migratory; growth-coupled essentiality asks which genes a tip cell needs in order to divide, which may be the wrong question for that state and may contribute to the non-recovery of PFKFB3 and GLUT1. A task-based essentiality analysis, in which a gene counts as essential if it makes any curated task infeasible, is the appropriate test and has not been performed here. Eighth, loop correction of flux variability ranges used a CycleFreeFlux-style procedure, which gives a conservative estimate of the loop-free range; exact loopless variability analysis by mixed-integer programming was not run on the full network. Finally, the most direct validation, namely genetic perturbation in zebrafish scored against intersegmental vessel sprouting with selective sparing of established vasculature, remains to be performed. Two experiments follow from this work: imaging of sprouting in the existing *slc25a20* mutant [65], which tests the candidate this screen would have reported without its checks, and perturbation of *phgdh*, which tests whether the predicted biosynthetic capacity of sprouting vessels depends on serine synthesis. A negative result in either would be informative, and we would report it.

## 5. Conclusions

We have presented FluxSprout, a set of endothelial state-specific genome-scale metabolic models of the sprouting zebrafish trunk, and used it to develop four checks on constraint-based vulnerability screening. A reaction by reaction audit showed that four inorganic pyrophosphatase reactions carry the oxidative phosphorylation subsystem label in the reference reconstruction; the apparent pathway dependence this produced arose only with every exchange open, where growth is set by arbitrary flux bounds, and under a defined medium the dependence on the corrected pathway is governed by the objective and the maintenance demand. An audit of internal cycles showed that a fifth of the network can carry flux without net substrate conversion, including every candidate of the conventional screen, so flux ranges and ratios for those reactions require loop correction. A two-level test showed that the tip-like and stalk-like intersegmental vessel populations differ mainly in cell-cycle phase, that glycolytic elevation distinguishes sprouting from established vessels at the marker level but not tip from stalk, and that the one flux difference between sprouting and established vessels robust to the capacity of the expression constraints is a higher predicted growth capacity, which serine and nucleotide synthesis follow in fixed proportion. An ensemble of four extraction settings, evaluated against every non-endothelial lineage, left no selective candidate, and traced the strongest single-pipeline call to one reaction omitted because one gene fell just below a threshold. Each of the four checks changed a conclusion we would otherwise have reported, which is the reason we present them as the subject of the paper rather than as caveats to it. None is specific to angiogenesis, and we offer them together as a template for any system in which a genome-scale model and single-cell data are both available.

## Acknowledgments

M.R.Z. is supported by NSF FDT-BioTech, Award Number #2533961 to R.C. which constituted the primary support for this work. Some computational resources used in this work were obtained through NSF 22-599, EPSCoR RII Track-1, Award Number #2242763 to R.C.

## Data and Code Availability

The single-cell RNA sequencing data are available under GEO accessions GSM6138416 and GSM6138417, and Zebrafish-GEM and Human-GEM from their repositories. The analysis notebook and code, the reaction lists of the 108 context-specific models (27 groups in each of four ensemble members), the medium definitions, the curated pathway definitions, the marker gene panels and all derived tables are deposited at https://github.com/ChowdhuryRatul/FluxSprout. Model quality metrics are given in Table S5. Supplementary Notes S1–S5, Figures S1–S4, Tables S1–S5 and Supplementary Data S1 accompany this article.

## References

[1] M. Potente, H. Gerhardt, and P. Carmeliet, “Basic and therapeutic aspects of angiogenesis,” Cell, vol. 146, no. 6, pp. 873–887, Sep. 2011, doi: 10.1016/J.CELL.2011.08.039.

[2] P. Carmeliet and R. K. Jain, “Molecular mechanisms and clinical applications of angiogenesis,” Nature, vol. 473, no. 7347, pp. 298–307, May 2011, doi: 10.1038/NATURE10144.

[3] H. Gerhardt et al., “VEGF guides angiogenic sprouting utilizing endothelial tip cell filopodia,” J. Cell Biol., vol. 161, no. 6, pp. 1163–1177, Jun. 2003, doi: 10.1083/JCB.200302047.

[4] M. Hellström et al., “Dll4 signalling through Notch1 regulates formation of tip cells during angiogenesis,” Nature, vol. 445, no. 7129, pp. 776–780, Feb. 2007, doi: 10.1038/NATURE05571.

[5] L. Jakobsson et al., “Endothelial cells dynamically compete for the tip cell position during angiogenic sprouting,” Nat. Cell Biol., vol. 12, no. 10, pp. 943–953, Oct. 2010, doi: 10.1038/NCB2103.

[6] K. De Bock et al., “Role of PFKFB3-driven glycolysis in vessel sprouting,” Cell, vol. 154, no. 3, Aug. 2013, doi: 10.1016/J.CELL.2013.06.037.

[7] S. Schoors et al., “Partial and transient reduction of glycolysis by PFKFB3 blockade reduces pathological angiogenesis,” Cell Metab., vol. 19, no. 1, pp. 37–48, Jan. 2014, doi: 10.1016/J.CMET.2013.11.008.

[8] A. R. Cantelmo et al., “Inhibition of the Glycolytic Activator PFKFB3 in Endothelium Induces Tumor Vessel Normalization, Impairs Metastasis, and Improves Chemotherapy,” Cancer Cell, vol. 30, no. 6, pp. 968–985, Dec. 2016, doi: 10.1016/J.CCELL.2016.10.006.

[9] B. Cruys et al., “Glycolytic regulation of cell rearrangement in angiogenesis,” Nature Communications 2016 7:1, vol. 7, no. 1, pp. 12240-, Jul. 2016, doi: 10.1038/ncomms12240.

[10] S. Schoors et al., “Fatty acid carbon is essential for dNTP synthesis in endothelial cells,” Nature, vol. 520, no. 7546, pp. 192–197, Apr. 2015, doi: 10.1038/NATURE14362.

[11] J. Kalucka et al., “Quiescent Endothelial Cells Upregulate Fatty Acid β-Oxidation for Vasculoprotection via Redox Homeostasis,” Cell Metab., vol. 28, no. 6, pp. 881–894.e13, Dec. 2018, doi: 10.1016/J.CMET.2018.07.016.

[12] B. W. Wong et al., “The role of fatty acid β-oxidation in lymphangiogenesis,” Nature, vol. 542, no. 7639, pp. 49–54, Feb. 2017, doi: 10.1038/NATURE21028.

[13] S. Vandekeere et al., “Serine Synthesis via PHGDH Is Essential for Heme Production in Endothelial Cells,” Cell Metab., vol. 28, no. 4, pp. 573–587.e13, Oct. 2018, doi: 10.1016/J.CMET.2018.06.009.

[14] H. Huang et al., “Role of glutamine and interlinked asparagine metabolism in vessel formation,” EMBO J., vol. 36, no. 16, pp. 2334–2352, Aug. 2017, doi: 10.15252/EMBJ.201695518.

[15] B. Kim, J. Li, C. Jang, and Z. Arany, “Glutamine fuels proliferation but not migration of endothelial cells,” EMBO J., vol. 36, no. 16, pp. 2321–2333, Aug. 2017, doi: 10.15252/EMBJ.201796436.

[16] U. Bruning et al., “Impairment of Angiogenesis by Fatty Acid Synthase Inhibition Involves mTOR Malonylation,” Cell Metab., vol. 28, no. 6, pp. 866–880.e15, Dec. 2018, doi: 10.1016/J.CMET.2018.07.019.

[17] G. Eelen, P. de Zeeuw, L. Treps, U. Harjes, B. W. Wong, and P. Carmeliet, “Endothelial Cell Metabolism,” Physiol. Rev., vol. 98, no. 1, pp. 3–58, Jan. 2018, doi: 10.1152/PHYSREV.00001.2017.

[18] X. Li, X. Sun, and P. Carmeliet, “Hallmarks of Endothelial Cell Metabolism in Health and Disease,” Cell Metab., vol. 30, no. 3, pp. 414–433, Sep. 2019, doi: 10.1016/J.CMET.2019.08.011.

[19] G. Bergers and D. Hanahan, “Modes of resistance to anti-angiogenic therapy,” Nat. Rev. Cancer, vol. 8, no. 8, pp. 592–603, Aug. 2008, doi: 10.1038/NRC2442.

[20] L. P. Diebold, H. J. Gil, P. Gao, C. A. Martinez, S. E. Weinberg, and N. S. Chandel, “Mitochondrial complex III is necessary for endothelial cell proliferation during angiogenesis,” Nat. Metab., vol. 1, no. 1, pp. 158–171, Jan. 2019, doi: 10.1038/S42255-018-0011-X.

[21] B. Yetkin-Arik et al., “The role of glycolysis and mitochondrial respiration in the formation and functioning of endothelial tip cells during angiogenesis,” Sci. Rep., vol. 9, no. 1, p. 12608, Dec. 2019, doi: 10.1038/S41598-019-48676-2.

[22] B. Yetkin-Arik et al., “Endothelial tip cells in vitro are less glycolytic and have a more flexible response to metabolic stress than non-tip cells,” Sci. Rep., vol. 9, no. 1, Dec. 2019, doi: 10.1038/S41598-019-46503-2.

[23] J. Kalucka et al., “Single-Cell Transcriptome Atlas of Murine Endothelial Cells,” Cell, vol. 180, no. 4, pp. 764–779.e20, Feb. 2020, doi: 10.1016/J.CELL.2020.01.015.

[24] J. Goveia et al., “An Integrated Gene Expression Landscape Profiling Approach to Identify Lung Tumor Endothelial Cell Heterogeneity and Angiogenic Candidates,” Cancer Cell, vol. 37, no. 1, pp. 21–36.e13, Jan. 2020, doi: 10.1016/J.CCELL.2019.12.001.

[25] K. Rohlenova et al., “Single-Cell RNA Sequencing Maps Endothelial Metabolic Plasticity in Pathological Angiogenesis,” Cell Metab., vol. 31, no. 4, pp. 862–877.e14, Apr. 2020, doi: 10.1016/J.CMET.2020.03.009.

[26] S. Gurung, N. K. Restrepo, B. Chestnut, L. Klimkaite, and S. Sumanas, “Single-cell transcriptomic analysis of vascular endothelial cells in zebrafish embryos,” Scientific Reports 2022 12:1, vol. 12, no. 1, pp. 13065-, Jul. 2022, doi: 10.1038/s41598-022-17127-w.

[27] H. B. Kwon et al., “In vivo modulation of endothelial polarization by Apelin receptor signalling,” Nat. Commun., vol. 7, Jun. 2016, doi: 10.1038/NCOMMS11805.

[28] F. Buettner et al., “Computational analysis of cell-to-cell heterogeneity in single-cell RNA-sequencing data reveals hidden subpopulations of cells,” Nat. Biotechnol., vol. 33, no. 2, pp. 155–160, 2015, doi: 10.1038/NBT.3102.

[29] N. D. Lawson and B. M. Weinstein, “In vivo imaging of embryonic vascular development using transgenic zebrafish,” Dev. Biol., vol. 248, no. 2, pp. 307–318, 2002, doi: 10.1006/dbio.2002.0711.

[30] S. Isogai, N. D. Lawson, S. Torrealday, M. Horiguchi, and B. M. Weinstein, “Angiogenic network formation in the developing vertebrate trunk,” Development, vol. 130, no. 21, pp. 5281–5290, Nov. 2003, doi: 10.1242/DEV.00733.

[31] A. F. Siekmann and N. D. Lawson, “Notch signalling limits angiogenic cell behaviour in developing zebrafish arteries,” Nature, vol. 445, no. 7129, pp. 781–784, Feb. 2007, doi: 10.1038/NATURE05577.

[32] J. D. Orth, I. Thiele, and B. O. Palsson, “What is flux balance analysis?,” Nat. Biotechnol., vol. 28, no. 3, pp. 245–248, Mar. 2010, doi: 10.1038/NBT.1614.

[33] C. Gu, G. B. Kim, W. J. Kim, H. U. Kim, and S. Y. Lee, “Current status and applications of genome-scale metabolic models,” Genome Biol., vol. 20, no. 1, Jun. 2019, doi: 10.1186/S13059-019-1730-3.

[34] A. Ebrahim, J. A. Lerman, B. O. Palsson, and D. R. Hyduke, “COBRApy: COnstraints-Based Reconstruction and Analysis for Python,” BMC Syst. Biol., vol. 7, Aug. 2013, doi: 10.1186/1752-0509-7-74.

[35] L. Heirendt et al., “Creation and analysis of biochemical constraint-based models using the COBRA Toolbox v.3.0,” Nat. Protoc., vol. 14, no. 3, pp. 639–702, Mar. 2019, doi: 10.1038/S41596-018-0098-2.

[36] C. Colijn et al., “Interpreting Expression Data with Metabolic Flux Models: Predicting Mycobacterium tuberculosis Mycolic Acid Production,” PLoS Comput. Biol., vol. 5, no. 8, p. e1000489, 2009, doi: 10.1371/JOURNAL.PCBI.1000489.

[37] N. Vlassis, M. P. Pacheco, and T. Sauter, “Fast Reconstruction of Compact Context-Specific Metabolic Network Models,” PLoS Comput. Biol., vol. 10, no. 1, p. e1003424, 2014, doi: 10.1371/JOURNAL.PCBI.1003424.

[38] R. Agren, S. Bordel, A. Mardinoglu, N. Pornputtapong, I. Nookaew, and J. Nielsen, “Reconstruction of Genome-Scale Active Metabolic Networks for 69 Human Cell Types and 16 Cancer Types Using INIT,” PLoS Comput. Biol., vol. 8, no. 5, p. e1002518, 2012, doi: 10.1371/JOURNAL.PCBI.1002518.

[39] S. A. Becker and B. O. Palsson, “Context-Specific Metabolic Networks Are Consistent with Experiments,” PLoS Comput. Biol., vol. 4, no. 5, p. e1000082, 2008, doi: 10.1371/JOURNAL.PCBI.1000082.

[40] S. Opdam, A. Richelle, B. Kellman, S. Li, D. C. Zielinski, and N. E. Lewis, “A Systematic Evaluation of Methods for Tailoring Genome-Scale Metabolic Models,” Cell Syst., vol. 4, no. 3, pp. 318–329.e6, Mar. 2017, doi: 10.1016/J.CELS.2017.01.010.

[41] D. Machado and M. Herrgård, “Systematic Evaluation of Methods for Integration of Transcriptomic Data into Constraint-Based Models of Metabolism,” PLoS Comput. Biol., vol. 10, no. 4, p. e1003580, 2014, doi: 10.1371/JOURNAL.PCBI.1003580.

[42] A. Richelle, C. Joshi, and N. E. Lewis, “Assessing key decisions for transcriptomic data integration in biochemical networks,” PLoS Comput. Biol., vol. 15, no. 7, p. e1007185, Jul. 2019, doi: 10.1371/JOURNAL.PCBI.1007185.

[43] J. Schellenberger, N. E. Lewis, and B. Palsson, “Elimination of Thermodynamically Infeasible Loops in Steady-State Metabolic Models,” Biophys. J., vol. 100, no. 3, pp. 544–553, Feb. 2011, doi: 10.1016/J.BPJ.2010.12.3707.

[44] A. A. Desouki, F. Jarre, G. Gelius-Dietrich, and M. J. Lercher, “CycleFreeFlux: efficient removal of thermodynamically infeasible loops from flux distributions,” Bioinformatics, vol. 31, no. 13, pp. 2159–2165, Jul. 2015, doi: 10.1093/BIOINFORMATICS/BTV096.

[45] M. R. Zargar, S. Malla, R. Saha, and R. Chowdhury, “Thermodynamic Quality Control Reshapes Spatially Resolved Metabolic Flux Predictions in Prostate Cancer,” bioRxiv, p. 2026.06.25.734531, Oct. 2026, doi: 10.64898/2026.06.25.734531.

[46] J. L. Robinson et al., “An atlas of human metabolism,” Sci. Signal., vol. 13, no. 624, Mar. 2020, doi: 10.1126/SCISIGNAL.AAZ1482/SUPPL_FILE/AAZ1482_SM.PDF.

[47] H. Wang et al., “Genome-scale metabolic network reconstruction of model animals as a platform for translational research,” Proc. Natl. Acad. Sci. U. S. A., vol. 118, no. 30, p. e2102344118, Jul. 2021, doi: 10.1073/PNAS.2102344118/SUPPL_FILE/PNAS.2102344118.SD04.XLSX.

[48] F. Gatto, R. Ferreira, and J. Nielsen, “Pan-cancer analysis of the metabolic reaction network,” Metab. Eng., vol. 57, pp. 51–62, Jan. 2020, doi: 10.1016/J.YMBEN.2019.09.006.

[49] A. Wagner et al., “Metabolic modeling of single Th17 cells reveals regulators of autoimmunity,” Cell, vol. 184, no. 16, pp. 4168–4185.e21, Aug. 2021, doi: 10.1016/J.CELL.2021.05.045.

[50] N. Alghamdi et al., “A graph neural network model to estimate cell-wise metabolic flux using single-cell RNA-seq data,” Genome Res., vol. 31, no. 10, pp. 1867–1884, Oct. 2021, doi: 10.1101/GR.271205.120.

[51] M. S. Noor et al., “Next-generation metabolic models informed by biomolecular simulations,” Curr. Opin. Biotechnol., vol. 92, Apr. 2025, doi: 10.1016/J.COPBIO.2025.103259.

[52] F. A. Wolf, P. Angerer, and F. J. Theis, “SCANPY: large-scale single-cell gene expression data analysis,” Genome Biol., vol. 19, no. 1, Feb. 2018, doi: 10.1186/S13059-017-1382-0.

[53] S. L. Wolock, R. Lopez, and A. M. Klein, “Scrublet: Computational Identification of Cell Doublets in Single-Cell Transcriptomic Data,” Cell Syst., vol. 8, no. 4, pp. 281–291.e9, Apr. 2019, doi: 10.1016/J.CELS.2018.11.005.

[54] T. Stuart et al., “Comprehensive Integration of Single-Cell Data,” Cell, vol. 177, no. 7, pp. 1888–1902.e21, Jun. 2019, doi: 10.1016/J.CELL.2019.05.031.

[55] I. Korsunsky et al., “Fast, sensitive and accurate integration of single-cell data with Harmony,” Nat. Methods, vol. 16, no. 12, pp. 1289–1296, Dec. 2019, doi: 10.1038/S41592-019-0619-0.

[56] V. A. Traag, L. Waltman, and N. J. van Eck, “From Louvain to Leiden: guaranteeing well-connected communities,” Scientific Reports 2019 9:1, vol. 9, no. 1, pp. 5233-, Mar. 2019, doi: 10.1038/s41598-019-41695-z.

[57] I. Tirosh et al., “Dissecting the multicellular ecosystem of metastatic melanoma by single-cell RNA-seq,” Science (1979)., vol. 352, no. 6282, pp. 189–196, Apr. 2016, doi: 10.1126/SCIENCE.AAD0501/SUPPL_FILE/TIROSH.SM.PDF.

[58] N. E. Lewis et al., “Omic data from evolved E. coli are consistent with computed optimal growth from genome-scale models,” Mol. Syst. Biol., vol. 6, 2010, doi: 10.1038/MSB.2010.47.

[59] R. Mahadevan and C. H. Schilling, “The effects of alternate optimal solutions in constraint-based genome-scale metabolic models,” Metab. Eng., vol. 5, no. 4, pp. 264–276, 2003, doi: 10.1016/j.ymben.2003.09.002.

[60] Y. Baran et al., “MetaCell: analysis of single-cell RNA-seq data using K-nn graph partitions,” Genome Biol., vol. 20, no. 1, Oct. 2019, doi: 10.1186/S13059-019-1812-2.

[61] Y. Benjamini and Y. Hochberg, “Controlling the False Discovery Rate: A Practical and Powerful Approach to Multiple Testing,” J. R. Stat. Soc. Series B Stat. Methodol., vol. 57, no. 1, pp. 289–300, Jan. 1995, doi: 10.1111/J.2517-6161.1995.TB02031.X.

[62] N. Cliff, “Dominance statistics: Ordinal analyses to answer ordinal questions,” Psychol. Bull., vol. 114, no. 3, pp. 494–509, 1993, doi: 10.1037/0033-2909.114.3.494.

[63] Y. Hu et al., “An integrative approach to ortholog prediction for disease-focused and other functional studies,” BMC Bioinformatics, vol. 12, Aug. 2011, doi: 10.1186/1471-2105-12-357.

[64] Q. Huangfu and J. A. J. Hall, “Parallelizing the dual revised simplex method,” Math. Program. Comput., vol. 10, no. 1, pp. 119–142, Mar. 2018, doi: 10.1007/S12532-017-0130-5/FIGURES/5.

[65] R. Hishida, K. Ishiguro, T. Yamanaka, S. Toyokuni, and H. Matsui, “Homozygous slc25a20 zebrafish mutant reveals insights into carnitine-acylcarnitine translocase deficiency pathogenesis,” Mol. Genet. Metab. Rep., vol. 41, p. 101165, Dec. 2024, doi: 10.1016/J.YMGMR.2024.101165.

[66] C. Indiveri et al., “The mitochondrial carnitine/acylcarnitine carrier: function, structure and physiopathology,” Mol. Aspects Med., vol. 32, no. 4–6, pp. 223–233, Aug. 2011, doi: 10.1016/J.MAM.2011.10.008.

[67] A. Sur, Y. Wang, P. Capar, G. Margolin, M. K. Prochaska, and J. A. Farrell, “Single-cell analysis of shared signatures and transcriptional diversity during zebrafish development,” Dev. Cell, vol. 58, no. 24, pp. 3028–3047.e12, Dec. 2023, doi: 10.1016/J.DEVCEL.2023.11.001.

[68] M. Lange et al., “A multimodal zebrafish developmental atlas reveals the state-transition dynamics of late-vertebrate pluripotent axial progenitors,” Cell, vol. 187, no. 23, pp. 6742–6759.e17, Nov. 2024, doi: 10.1016/J.CELL.2024.09.047.

[69] H. Zur, E. Ruppin, and T. Shlomi, “iMAT: an integrative metabolic analysis tool,” Bioinformatics, vol. 26, no. 24, pp. 3140–3142, Dec. 2010, doi: 10.1093/BIOINFORMATICS/BTQ602.

