## Supplementary Information for "FluxSprout: Single-cell metabolic modeling of vascular sprouting in zebrafish"

**Contents**

Supplementary Note S1. The conventional single-pipeline screen

Supplementary Note S2. Linear programming engine and its validation

Supplementary Note S3. Context-specific extraction: FASTCORE, GIMME and the closure step

Supplementary Note S4. Loop audit and loop correction

Supplementary Note S5. Metacell models and flux read-outs

Supplementary Figures S1–S4

Supplementary Tables S1–S5

Supplementary Data S1 (FluxSprout_Supplementary_Data.xlsx)

Code and data

References in square brackets refer to the reference list of the main text.

#### Supplementary Notes

##### Note S1. The conventional single-pipeline screen

To show what each check of the main text changes, the vulnerability screen was also run in the way such screens are commonly performed. We refer to this run as the conventional screen. It differs from the analysis of the main text in four respects, each of which corresponds to one of the four checks.

(i) Pathway annotation. Oxidative phosphorylation was defined by its subsystem label, that is, by all ten reactions carrying the label, including the four inorganic pyrophosphatases (Section 2.7, Table 3). Pathway titration was performed on the four-compartment models with every exchange reaction open. (ii) Loops. Flux variability ranges and flux ratios were reported without loop correction. (iii) Expression constraints. E-Flux bounds were scaled from the default bound of ±1000 after normalizing each reaction score to the maximum across reactions (the "conv." setting of Fig. 5A), and a single pseudobulk model was built per state or compartment, with no replicate units. (iv) Extraction and specificity. One set of thirteen state models was extracted with FASTCORE at a single, default local-T2 threshold, without a maintenance demand, and a candidate was called if it inhibited sprouting growth by at least 0.10 while sparing the erythroid and skeletal muscle models (the two-lineage rule of Section 2.10). Every gene with a GPR association to one of the ten curated pathways (2,097 genes) was screened.

The conventional screen nominated SLC25A20, SLC25A36 and IDH2 as vulnerabilities selective for sprouting endothelium, and its titration suggested that about 64% of growth depended on oxidative phosphorylation. In the main text these outputs serve only as a reference against which the four checks are evaluated (Sections 3.4, 3.5 and 3.7, Fig. 4E, Fig. 6, Table 4 and Table 5); none of them is reported as a finding of this study.

##### Note S2. Linear programming engine and its validation

Flux balance analysis (FBA), parsimonious FBA, flux variability analysis (FVA), loop correction, gene deletion, FASTCORE and GIMME were implemented on a purpose-written interface to the HiGHS solver [64] (highspy 1.15), which holds the stoichiometric matrix in sparse form and reuses the solver basis between related problems. Gene deletions are applied by evaluating each reaction's GPR rule with the deleted gene set to zero (AND, minimum; OR, maximum) and closing every reaction whose rule evaluates to zero. COBRApy 0.32 [34] was used for model input, mass and charge balance checks and the reference comparisons. On the *E. coli* core model the interface reproduces COBRApy results to within $10^{-11}$ for FBA, FVA and single-gene deletion, and identifies the same blocked reactions. A single deletion on the 9,480-reaction reference network takes about 0.03 s with warm starts, which made the full screen (825 genes in 27 groups and four ensemble members, about 89,000 deletions) feasible.

##### Note S3. Context-specific extraction: FASTCORE, GIMME and the closure step

Activity calls. Gene activity was called per group by local-T2 thresholding [42] of the log1p-transformed mean CP10k expression: a gene is active in a group if its expression exceeds the global upper threshold, or exceeds the global lower threshold and its own mean across the 27 groups. The global thresholds are percentiles of all gene–group values: 15th/65th (loose), 25th/75th (default) and 40th/85th (strict). A reaction is active if its GPR rule evaluates to active.

FASTCORE. FASTCORE [37] was reimplemented with its LP7 and LP10 subproblems. The core comprised the active reactions, the biomass reaction MAR13082 and the four biomass pool reactions

(MAR10062–MAR10065). Exchange reactions were excluded from the core but retained in every model, together with the biomass, pool and maintenance reactions. Extraction was run with the maintenance demand released, because with a forced maintenance flux the LP7 constraint cannot be satisfied when uptake exchanges are in the core. Flux consistency of the reference network (FASTCC) was established by batched LP7 problems that maximize the number of reactions carrying at least $10^{-5}$, forward and then backward, followed by random-objective probes until five successive probes found no new reaction.

Closure step. The support threshold of FASTCORE drops micronutrient routes whose fluxes are of the order of $10^{-6}$ of the biomass flux, which leaves the biomass reaction unable to carry flux. After extraction, the minimal-flux set of reference reactions outside the extracted model that allows biomass at 10% of the reference optimum, with the maintenance demand restored (NGAM = 1), was added to each model.

GIMME. GIMME [39] minimized $\Sigma_j (\theta - s_j)|v_j|$ over the reactions whose GPR score $s_j$ falls below $\theta$, subject to growth of at least 90% of the reference optimum, with $\theta$ the default local-T2 upper threshold. The extracted model comprised the reactions carrying flux above $10^{-7}$ in that solution, every reaction scoring at or above $\theta$, and the exchange, biomass, pool and maintenance reactions. Model sizes and the agreement between members are given in Table S3 and Fig. S4A–B.

##### Note S4. Loop audit and loop correction

Audit. All exchange reactions, the three biomass reactions and the maintenance reaction were closed in the flux-consistent network (11,356 reactions with every exchange open), and the reactions that could still carry flux were identified with the procedure used for flux consistency (Note S3). Any such reaction can carry flux only around an internal cycle. The 2,217 loop-capable reactions are listed in Supplementary Data S1 (sheet D9).

Nearest loop-free flux. For a flux vector $v$, the nearest loop-free flux was obtained as in CycleFreeFlux [44]: exchange fluxes are fixed at their values in $v$, each internal reaction is confined to the interval between zero and its value in $v$, and the total absolute internal flux is minimized. The L1 distance between $v$ and this solution measures loop flux; it was below $10^{-7}$ for every parsimonious solution in this study (Fig. 5E).

Loop-corrected FVA. For each extreme of an FVA range, the optimal flux vector was replaced by its nearest loop-free flux. If the target reaction kept its value, the extreme was accepted; otherwise the target was re-optimized with every reaction constrained to the sign pattern of the loop-free solution. Each reported bound is therefore attained by a loop-free flux distribution, so the corrected range lies inside the true loop-free range and is a conservative estimate of it. A reaction is flagged as loop-dependent when its corrected range is narrower than the uncorrected one (Table 6). Exact loopless FVA by mixed-integer programming [43] was not run on the full network.

##### Note S5. Metacell models and flux read-outs

Within each state and lineage, cells were partitioned by k-means on the first 30 principal components into $k = \max(3, n/20)$ groups, and groups of fewer than five cells were discarded, giving 384 metacells. E-Flux models were built for the 325 metacells of the fourteen endothelial and hemogenic states and five non-endothelial lineages (erythroid, macrophage, neural progenitor, fibroblast and skeletal muscle), from the mean CP10k expression of each metacell, at U = 50 and U = 100 (Table S3). Each model was solved by parsimonious FBA and checked for loops (Note S4). The read-outs and the reactions that define them are listed below.

| Read-out | Reactions | Definition |
|---|---|---|
| Predicted growth | MAR13082 | Growth relative to the unconstrained reference network |
| Glycolytic share of ATP production | MAR04368 (PGK), MAR04358 (PK), MAR06916 (ATP synthase) | (PGK + PK) / (PGK + PK + ATP synthase), forward fluxes |
| Lactate per glucose | MAR09135, MAR09034 | Lactate secretion / glucose uptake |
| $O_2$ uptake | MAR09048 | Oxygen exchange flux |
| PHGDH flux | MAR03839 | Phosphoglycerate dehydrogenase flux |
| Carnitine–acylcarnitine carrier flux | MAR02629 | Mitochondrial carnitine–acylcarnitine exchange (SLC25A20) |
| De novo purine synthesis per unit growth | MAR04406 (PPAT) | PPAT flux / growth |
| De novo pyrimidine synthesis per unit growth | MAR04034 (CAD, CPS II) | CPS II flux / growth |

In every sprouting and established-vessel metacell, PPAT flux was 0.102 times growth and PHGDH flux 0.0012 times growth, so differences between groups in these two read-outs are differences in predicted growth (Section 3.6, Table 5).

### Supplementary Figures

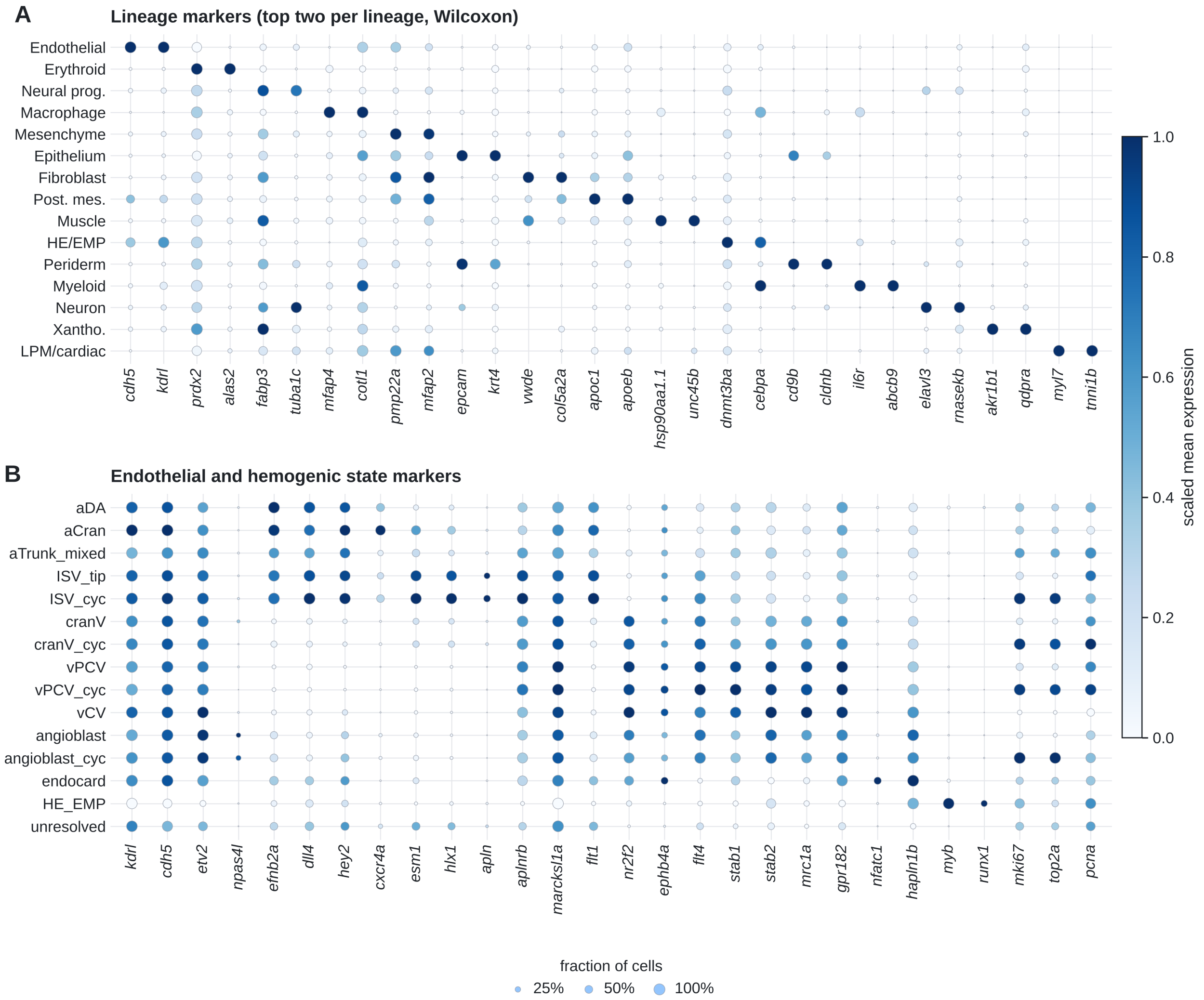


**Fig. S1. Marker genes of the lineages and of the endothelial and hemogenic states.** (A) The two top-ranked marker genes of each lineage (Wilcoxon rank-sum test against all other annotated cells; ribosomal, mitochondrial, hemoglobin and unannotated genes excluded). (B) Curated markers of the fourteen endothelial and hemogenic states and the unresolved cluster: pan-endothelial (*kdrl*, *cdh5*, *etv2*), angioblast (*npas4l*), arterial (*efnb2a*, *dll4*, *hey2*, *cxcr4a*), sprouting intersegmental vessel (*esm1*, *hlx1*, *apln*, *aplnrb*, *marcksl1a*, *flt1*), venous (*nr2f2*, *ephb4a*, *flt4*, *stab1*, *stab2*, *mrc1a*, *gpr182*), endocardial (*nfatc1*, *hapln1b*), hemogenic (*myb*, *runx1*) and proliferation (*mki67*, *top2a*, *pcna*). Dot size, fraction of cells in which the gene is detected; color, mean expression scaled from 0 to 1 per gene across groups.

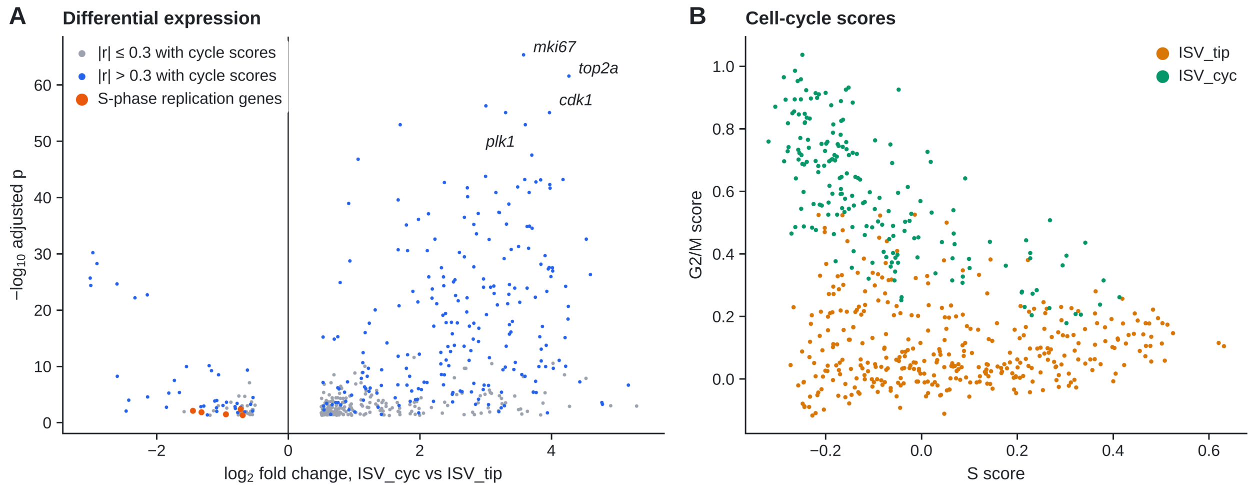


**Fig. S2. The difference between ISV_cyc and ISV_tip is a cell-cycle difference.** (A) Genes differentially expressed between ISV_cyc and ISV_tip (Wilcoxon rank-sum test; $q < 0.05$ and $|\log_2 \text{fold change}| > 0.5$; 449 up and 72 down in ISV_cyc), colored by whether their expression correlates with the S or G2/M score across endothelial cells ($|r| > 0.3$). Orange, the S-phase replication genes *mcm7*, *rfc4*, *pold1*, *fen1* and *gins2*, which are lower in ISV_cyc. (B) S and G2/M scores of the 547 ISV cells.

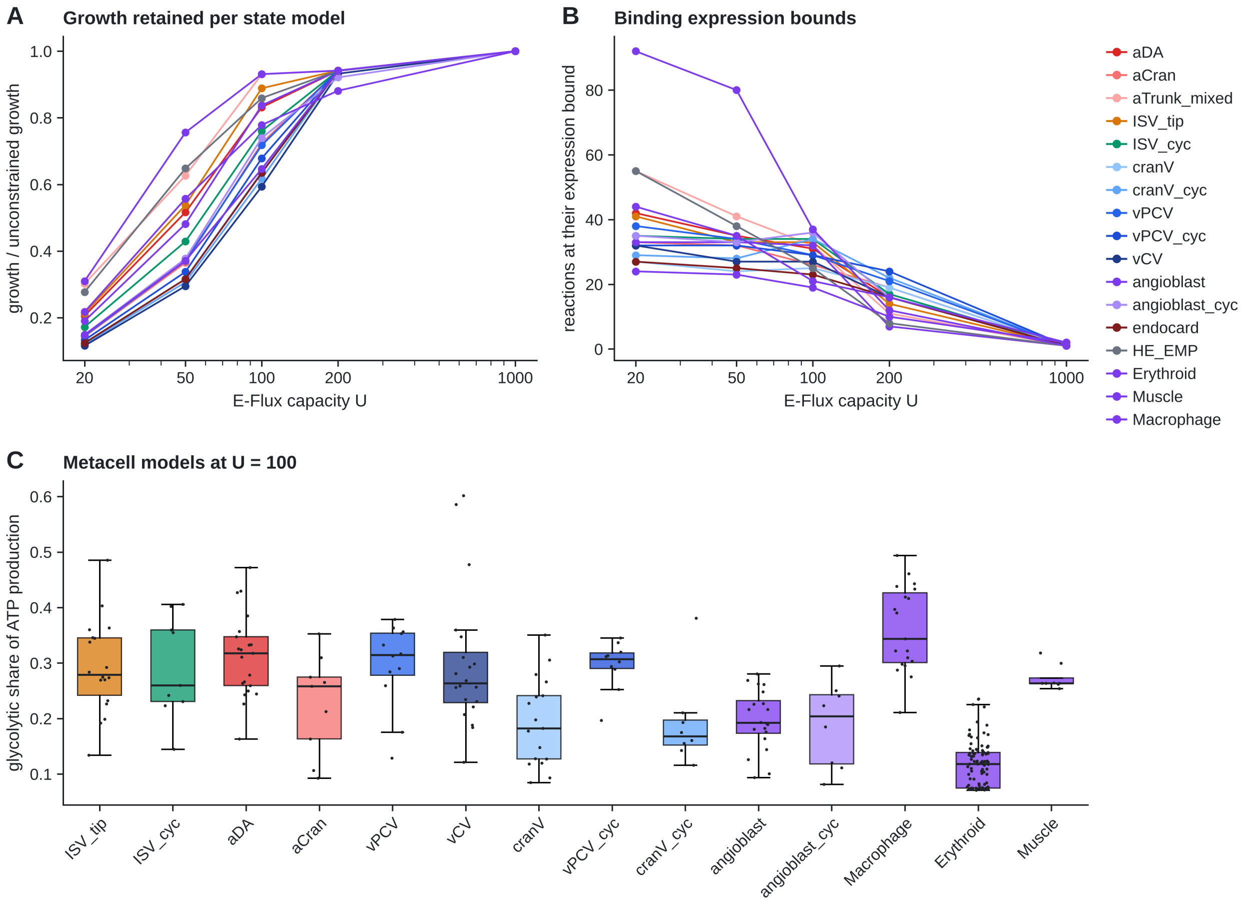


**Fig. S3. Calibration of the E-Flux capacity.** (A) Growth of each pseudobulk state and lineage model relative to the unconstrained reference network, and (B) number of internal reactions at their expression bound, as the capacity U is varied (reaction scores normalized to their 95th percentile). (C) Glycolytic share of ATP production in the metacell models at U = 100, the sensitivity setting (compare Fig. 5C at U = 50). Boxes, interquartile range and median; points, metacells.

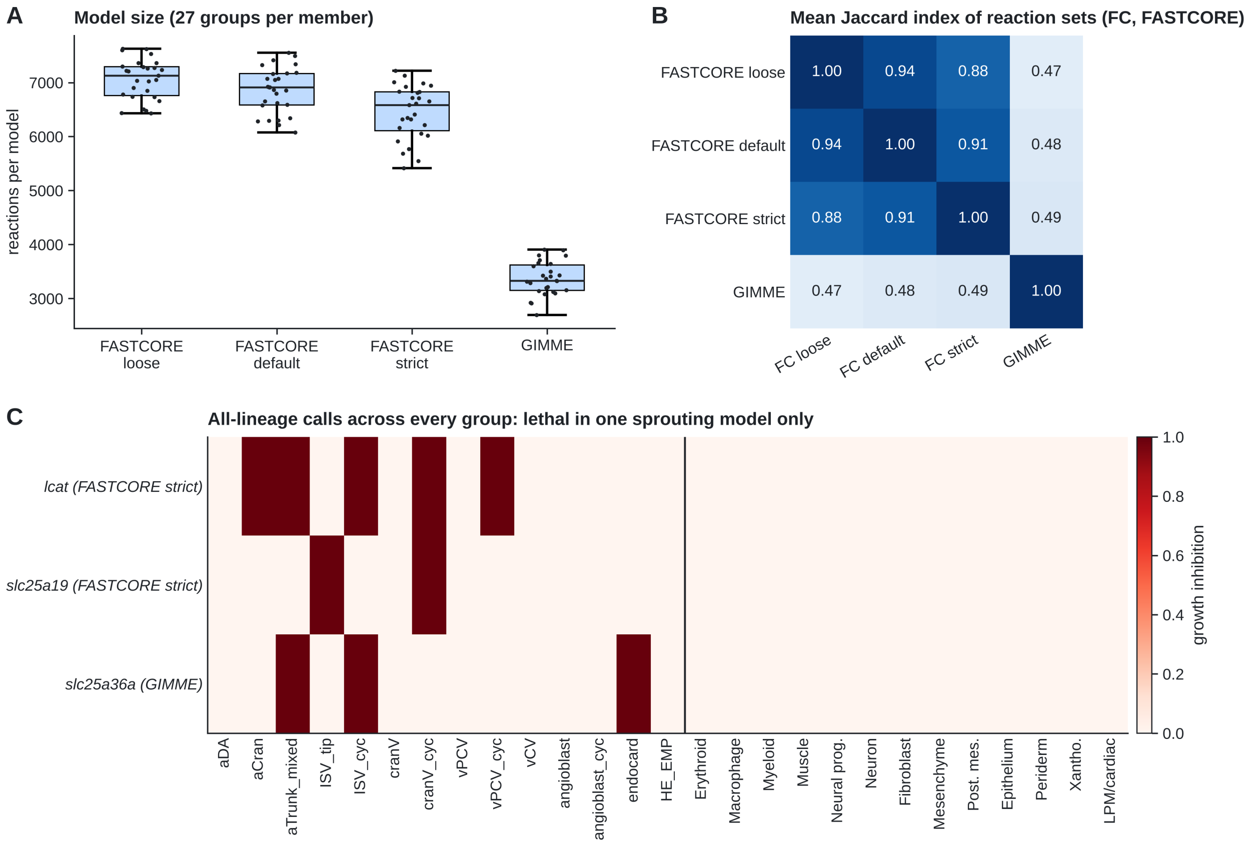


**Fig. S4. Structure of the four-member model ensemble and the three all-lineage calls.** (A) Number of reactions in the 27 group models of each ensemble member. (B) Mean Jaccard index between the reaction sets of the same group in two members (FC, FASTCORE). (C) Growth inhibition by the three genes called under the all-lineage rule, in every endothelial and hemogenic state (left of the vertical line) and every non-endothelial lineage (right), in the member that called each gene. Each call is lethal in exactly one of the two sprouting models and spares every non-endothelial lineage; the reactions responsible are given in Table 4 and Supplementary Data S1 (sheet D12).

#### Supplementary Tables

**Table S1. Quality control and cell numbers.** (a) Droplets per library before filtering and removed by each filter; the filters overlap, so the removed counts do not sum to the difference between raw and retained droplets. The upper-tail outlier filter (more than five median absolute deviations above the median total count) removed no droplet. (b) Annotated cells per lineage. (c) Endothelial and hemogenic cells per state.

### (a)

| Library | Raw | Low genes or UMIs | High mito. | MAD outlier | Doublet | Retained |
|---|---|---|---|---|---|---|
| $GFP^{+}$ $mCherry^{-}$ (GSM6138417) | 6,990 | 1,443 | 1,049 | 0 | 85 | 5,024 |
| $GFP^{+}$ $mCherry^{+}$ (GSM6138416) | 6,390 | 813 | 872 | 0 | 43 | 5,066 |

##### (b) and (c)

| Lineage | Cells | State | Cells |
|---|---|---|---|
| Endothelial | 3596 | vCV | 443 |
| Erythroid | 2536 | aDA | 430 |
| Neural prog. | 431 | angioblast | 428 |
| Macrophage | 391 | cranV | 364 |
| Mesenchyme | 320 | ISV_tip | 362 |
| Epithelium | 301 | vPCV | 292 |
| Fibroblast | 263 | unresolved | 222 |
| Post. mes. | 211 | vPCV_cyc | 202 |
| Muscle | 170 | aCran | 193 |
| HE/EMP | 131 | ISV_cyc | 185 |
| Periderm | 128 | cranV_cyc | 180 |
| Myeloid | 119 | angioblast_cyc | 169 |
| Neuron | 69 | aTrunk_mixed | 110 |
| Xantho. | 24 | HE_EMP | 107 |
| LPM/cardiac | 19 | endocard | 40 |

**Table S2. Medium and exchange bounds.** Maximum uptake (mmol $gDW^{-1}$ $h^{-1}$) of each nutrient in the baseline embryonic medium and in each sensitivity medium; all secretions are open. Only the bounds named in Section 2.10 differ between media. Vitamin B12 (aquacob(III)alamin) and γ-tocopherol are required for the cofactor pool of the biomass reaction on a defined medium.

| Nutrient | Exchange | Metabolite | Baseline | Hypoxia | Ischemia | Hyperglycemia | +Carnitine | Low amino acids |
|---|---|---|---|---|---|---|---|---|
| glucose | MAR09034 | glucose | 10 | 30 | 2 | 40 | 10 | 10 |
| O2 | MAR09048 | O2 | 20 | 1 | 0.8 | 20 | 20 | 20 |
| alanine | MAR09061 | alanine | 1 | 1 | 1 | 1 | 1 | 0.1 |
| arginine | MAR09066 | arginine | 1 | 1 | 1 | 1 | 1 | 0.1 |
| asparagine | MAR09062 | asparagine | 1 | 1 | 1 | 1 | 1 | 0.1 |
| aspartate | MAR09070 | aspartate | 1 | 1 | 1 | 1 | 1 | 0.1 |
| cysteine | MAR09065 | cysteine | 1 | 1 | 1 | 1 | 1 | 0.1 |
| glutamine | MAR09063 | glutamine | 10 | 20 | 2 | 10 | 10 | 10 |
| glutamate | MAR09071 | glutamate | 1 | 1 | 1 | 1 | 1 | 0.1 |
| glycine | MAR09067 | glycine | 1 | 1 | 1 | 1 | 1 | 0.1 |
| histidine | MAR09038 | histidine | 1 | 1 | 1 | 1 | 1 | 0.1 |
| isoleucine | MAR09039 | isoleucine | 1 | 1 | 1 | 1 | 1 | 0.1 |
| leucine | MAR09040 | leucine | 1 | 1 | 1 | 1 | 1 | 0.1 |
| lysine | MAR09041 | lysine | 1 | 1 | 1 | 1 | 1 | 0.1 |
| methionine | MAR09042 | methionine | 1 | 1 | 1 | 1 | 1 | 0.1 |
| phenylalanine | MAR09043 | phenylalanine | 1 | 1 | 1 | 1 | 1 | 0.1 |
| proline | MAR09068 | proline | 1 | 1 | 1 | 1 | 1 | 0.1 |
| serine | MAR09069 | serine | 1 | 1 | 1 | 1 | 1 | 0.1 |
| threonine | MAR09044 | threonine | 1 | 1 | 1 | 1 | 1 | 0.1 |
| tryptophan | MAR09045 | tryptophan | 1 | 1 | 1 | 1 | 1 | 0.1 |
| tyrosine | MAR09064 | tyrosine | 1 | 1 | 1 | 1 | 1 | 0.1 |
| valine | MAR09046 | valine | 1 | 1 | 1 | 1 | 1 | 0.1 |
| palmitate | MAR00611 | palmitate | 1 | 1 | 1 | 1 | 1 | 1 |
| oleate | MAR00650 | oleate | 1 | 1 | 1 | 1 | 1 | 1 |
| linoleate | MAR09035 | linoleate | 1 | 1 | 1 | 1 | 1 | 1 |
| linolenate | MAR09036 | linolenate | 1 | 1 | 1 | 1 | 1 | 1 |
| DHA | MAR00573 | DHA | 1 | 1 | 1 | 1 | 1 | 1 |
| arachidonate | MAR00568 | arachidonate | 1 | 1 | 1 | 1 | 1 | 1 |
| cholesterol | MAR09285 | cholesterol | 1 | 1 | 1 | 1 | 1 | 1 |
| choline | MAR09083 | choline | 1 | 1 | 1 | 1 | 1 | 1 |
| ethanolamine | MAR09084 | ethanolamine | 1 | 1 | 1 | 1 | 1 | 1 |
| folate | MAR09146 | folate | 0.1 | 0.1 | 0.1 | 0.1 | 0.1 | 0.1 |
| thiamin | MAR09159 | thiamin | 0.1 | 0.1 | 0.1 | 0.1 | 0.1 | 0.1 |
| biotin | MAR09109 | biotin | 0.1 | 0.1 | 0.1 | 0.1 | 0.1 | 0.1 |
| riboflavin | MAR09143 | riboflavin | 0.1 | 0.1 | 0.1 | 0.1 | 0.1 | 0.1 |
| nicotinamide | MAR09378 | nicotinamide | 0.1 | 0.1 | 0.1 | 0.1 | 0.1 | 0.1 |
| pyridoxine | MAR09144 | pyridoxine | 0.1 | 0.1 | 0.1 | 0.1 | 0.1 | 0.1 |
| pantothenate | MAR09145 | pantothenate | 0.1 | 0.1 | 0.1 | 0.1 | 0.1 | 0.1 |
| inositol | MAR09361 | inositol | 0.1 | 0.1 | 0.1 | 0.1 | 0.1 | 0.1 |
| retinol | MAR09147 | retinol | 0.1 | 0.1 | 0.1 | 0.1 | 0.1 | 0.1 |
| lipoic acid | MAR09167 | lipoic acid | 0.1 | 0.1 | 0.1 | 0.1 | 0.1 | 0.1 |
| ascorbate | MAR09158 | ascorbate | 0.1 | 0.1 | 0.1 | 0.1 | 0.1 | 0.1 |
| alpha-tocopherol | MAR09151 | alpha-tocopherol | 0.1 | 0.1 | 0.1 | 0.1 | 0.1 | 0.1 |
| Pi | MAR09072 | Pi | open | open | open | open | open | open |
| Fe2+ | MAR09076 | Fe2+ | open | open | open | open | open | open |
| sulfate | MAR09074 | sulfate | open | open | open | open | open | open |
| H2O | MAR09047 | H2O | open | open | open | open | open | open |
| H+ | MAR09079 | H+ | open | open | open | open | open | open |
| gamma-tocopherol | MAR09153 | gamma-tocopherol | 0.1 | 0.1 | 0.1 | 0.1 | 0.1 | 0.1 |
| B12 | MAR09269 | aquacob(III)alamin | 0.1 | 0.1 | 0.1 | 0.1 | 0.1 | 0.1 |
| carnitine | MAR09292 | L-carnitine | 0 | 0 | 0 | 0 | 1 | 0 |

Open, no upper limit on uptake (bound of 1000).

**Table S3. Groups, compartments, replicate units and model sizes.** Every endothelial and hemogenic state and every non-endothelial lineage, with the compartment used for statistical testing (Section 2.2), the number of cells, the number of metacells for which an E-Flux model was built, and the number of reactions in the model of each ensemble member. Sprouting, ISV_tip and ISV_cyc; established, aDA, aCran, vPCV, vCV and cranV.

| Group | Type | Compartment | Cells | Metacells modeled | FASTCORE loose | FASTCORE default | FASTCORE strict | GIMME |
|---|---|---|---|---|---|---|---|---|
| aDA | EC or HE state | arterial; established | 430 | 21 | 7290 | 7160 | 6830 | 3122 |
| aCran | EC or HE state | arterial; established | 193 | 9 | 7365 | 7328 | 6943 | 3659 |
| aTrunk_mixed | EC or HE state | atlas only | 110 | 5 | 6900 | 6655 | 6346 | 3099 |
| ISV_tip | EC or HE state | tip; sprouting | 362 | 18 | 7131 | 6863 | 6057 | 3082 |
| ISV_cyc | EC or HE state | cycling; sprouting | 185 | 9 | 7273 | 6915 | 6611 | 2923 |
| cranV | EC or HE state | venous; established | 364 | 17 | 7220 | 6925 | 6717 | 3158 |
| cranV_cyc | EC or HE state | cycling | 180 | 8 | 7023 | 6796 | 6019 | 3141 |
| vPCV | EC or HE state | venous; established | 292 | 12 | 7033 | 6854 | 6653 | 3502 |
| vPCV_cyc | EC or HE state | cycling | 202 | 10 | 7233 | 7074 | 6321 | 3366 |
| vCV | EC or HE state | venous; established | 443 | 20 | 7620 | 7344 | 7130 | 3899 |
| angioblast | EC or HE state | atlas only | 428 | 20 | 7365 | 7073 | 6831 | 3803 |
| angioblast_cyc | EC or HE state | cycling | 169 | 8 | 7300 | 7170 | 6923 | 3313 |
| endocard | EC or HE state | atlas only | 40 | 2 | 7261 | 7182 | 6985 | 3711 |
| HE_EMP | EC or HE state | atlas only | 107 | 5 | 7601 | 7494 | 7218 | 3410 |
| Erythroid | lineage | control | 2536 | 101 | 6740 | 6076 | 5414 | 2695 |
| Macrophage | lineage | control | 391 | 19 | 7531 | 7414 | 7006 | 3600 |
| Myeloid | lineage | control | 119 | 0 | 6732 | 6591 | 6408 | 3428 |
| Muscle | lineage | control | 170 | 8 | 6658 | 6285 | 5546 | 2914 |
| Neural prog. | lineage | control | 431 | 21 | 6438 | 6298 | 5912 | 3212 |
| Neuron | lineage | control | 69 | 0 | 7051 | 6911 | 6587 | 3905 |
| Fibroblast | lineage | control | 263 | 12 | 6783 | 6580 | 6213 | 3430 |
| Mesenchyme | lineage | control | 320 | 0 | 6503 | 6213 | 5686 | 3333 |
| Post. mes. | lineage | control | 211 | 0 | 6432 | 6292 | 5769 | 3323 |
| Epithelium | lineage | control | 301 | 0 | 6849 | 6622 | 6322 | 3286 |
| Periderm | lineage | control | 128 | 0 | 7626 | 7551 | 6712 | 3640 |
| Xantho. | lineage | control | 24 | 0 | 7207 | 7051 | 6812 | 3798 |
| LPM/cardiac | lineage | control | 19 | 0 | 6478 | 6342 | 6159 | 3196 |

**Table S4. Marker gene panels and cell-cycle genes.** Genes of each curated pathway's Zebrafish-GEM reactions that are detected in at least 5% of endothelial cells, used for the per-cell pathway scores (the four pyrophosphatase genes are excluded from the oxidative phosphorylation panel), and the S and G2/M genes of Tirosh et al. [57] recovered as zebrafish orthologs (including a and b paralogs).

| Panel | Genes | Gene symbols |
|---|---|---|
| Glycolysis / gluconeogenesis | 42 | *acss2*, *adh5*, *adh8b*, *adhfe1*, *adpgk*, *akr1a1b*, *aldh2.1*, *aldh2.2*, *aldh3a2a*, *aldh3a2b*, *aldh7a1*, *aldh9a1a.1*, *aldoaa*, *aldob*, *aldocb*, *bpgm*, *dlat*, *eno1a*, *eno3*, *fbp1a*, *g6pc3*, *galm*, *gapdh*, *gapdhs*, *gpd2*, *gpia*, *hk1*, *hk2*, *ldha*, *ldhba*, *pdha1a*, *pdhb*, *pdhx*, *pfklb*, *pfkpa*, *pgam1a*, *pgam1b*, *pgk1*, *pgm1*, *pgm2*, *pkma*, *tpi1b* |
| TCA cycle | 33 | *aclya*, *aco1*, *aco2*, *aldh2.1*, *aldh2.2*, *aldh3a2a*, *aldh3a2b*, *aldh7a1*, *aldh9a1a.1*, *aldoaa*, *aldob*, *aldocb*, *cs*, *dlst*, *fh*, *idh1*, *idh2*, *idh3a*, *idh3b*, *idh3g*, *ldha*, *ldhba*, *mdh1aa*, *mdh2*, *ogdha*, *oxct1a*, *sdha*, *sdhb*, *sdhc*, *sdhdb*, *sucla2*, *suclg1*, *suclg2* |
| Oxidative phosphorylation (corrected) | 85 | *atp5f1b*, *atp5f1c*, *atp5f1d*, *atp5f1e*, *atp5fa1*, *atp5l*, *atp5mc1*, *atp5mc3a*, *atp5mc3b*, *atp5mea*, *atp5meb*, *atp5mf*, *atp5pb*, *atp5pd*, *atp5pf*, *atp5po*, *cox4i1*, *cox4i2*, *cox5aa*, *cox6b1*, *cox6b2*, *cox6c*, *cox7a2a*, *cox7b*, *cox7c*, *cox8a*, *cyc1*, *dhodh*, *etfa*, *etfb*, *etfdh*, *gpd2*, *mt-atp6*, *mt-co1*, *mt-co2*, *mt-co3*, *mt-cyb*, *mt-nd1*, *mt-nd2*, *mt-nd3*, *mt-nd4*, *mt-nd4l*, *mt-nd5*, *mt-nd6*, *ndufa1*, *ndufa10*, *ndufa11*, *ndufa12*, *ndufa2*, *ndufa3*, *ndufa5*, *ndufa6*, *ndufa7*, *ndufa8*, *ndufa9a*, *ndufab1b*, *ndufb10*, *ndufb11*, *ndufb2*, *ndufb3*, *ndufb4*, *ndufb5*, *ndufb6*, *ndufb7*, *ndufb8*, *ndufb9*, *ndufc2*, *ndufs2*, *ndufs3*, *ndufs4*, *ndufs5*, *ndufs6*, *ndufs7*, *ndufs8a*, *ndufv1*, *ndufv2*, *ndufv3*, *prodha*, *uqcr10*, *uqcrb*, *uqcrc1*, *uqcrc2b*, *uqcrfs1*, *uqcrh*, *uqcrq* |
| Mitochondrial fatty acid oxidation | 19 | *acaa1*, *acaa2*, *acad9*, *acadl*, *acadm*, *acadsb*, *acadvl*, *acox1*, *decr1*, *decr2*, *ech1*, *echs1*, *eci1*, *eci2*, *hadh*, *hadhaa*, *hadhab*, *hadhb*, *hsd17b10* |
| Nucleotide synthesis | 82 | *acad8*, *acad9*, *acadm*, *acadvl*, *acp2*, *acp6*, *ada2a*, *adcy6a*, *adcy7*, *adka*, *adsl*, *ak1*, *ak2*, *ak3*, *ak4*, *aprt*, *atic*, *cad*, *cant1a*, *cmpk*, *ctps1a*, *dck*, *dguok*, *dhodh*, *dpysl2b*, *dtymk*, *dut*, *enpp1*, *entpd1*, *entpd4*, *entpd5b*, *entpd6*, *entpd8*, *fam20b*, *gmpr2*, *gmps*, *guk1a*, *hk1*, *hk2*, *hprt1*, *impdh1b*, *itpa*, *nme2b.1*, *nme3*, *nme6*, *nme7*, *nol9*, *npr1a*, *nt5c1bb*, *nt5c2a*, *nt5c2b*, *nt5c3a*, *nudt14*, *nudt5*, *nudt9*, *paics*, *pde10a*, *pde4ba*, *pde4d*, *pde7a*, *pkma*, *pnp5a*, *ppat*, *prdx6*, *rpusd4*, *rrm1*, *rrm2*, *rrm2b*, *taf9*, *tk1*, *tk2*, *txn*, *txn2*, *txnrd3*, *tyms*, *uck1*, *uck2a*, *umps*, *upb1*, *uprt*, *zgc:136858*, *zgc:86811* |
| Serine / one-carbon metabolism | 30 | *acads*, *acadsb*, *alas1*, *aldh1l1*, *aldh7a1*, *amt*, *aoc2*, *ckba*, *dhfr*, *dldh*, *gcat*, *ggh*, *ggt5a*, *glyctk*, *gnmt*, *hyi*, *ivd*, *mthfd1b*, *mthfd2*, *mthfr*, *mthfs*, *phgdh*, *psat1*, *psph*, *shmt1*, *si:dkey-76k16.6*, *tdh*, *tha1*, *ubac2*, *vps29* |
| Mitochondrial carnitine shuttle | 4 | *cpt2*, *crata*, *crot*, *slc25a20* |
| Pentose phosphate pathway | 14 | *aldh3a2b*, *fbp1a*, *g6pd*, *h6pd*, *pfklb*, *pfkpa*, *pgd*, *pgls*, *pgm2*, *prps1b*, *rbks*, *rpe*, *rpia*, *tktb* |
| S phase (Tirosh et al., zebrafish orthologs) | 45 | *mcm5*, *pcna*, *tyms*, *fen1*, *mcm2*, *mcm4*, *rrm1*, *unga*, *gins2*, *mcm6*, *cdca7a*, *cdca7b*, *dtl*, *prim1*, *uhrf1*, *hells*, *rfc2*, *rpa2*, *nasp*, *rad51ap1*, *gmnn*, *wdr76*, *slbp*, *ccne2*, *ubr7*, *pold3*, *msh2*, *atad2*, *atad2b*, *rad51b*, *rad51*, *rrm2*, *rrm2b*, *cdc45*, *cdc6*, *exo1*, *tipin*, *dscc1*, *blm*, *casp8ap2*, *usp1*, *pola1*, *chaf1b*, *brip1*, *e2f8* |
| G2/M phase (Tirosh et al., zebrafish orthologs) | 50 | *hmgb2a*, *hmgb2b*, *cdk1*, *nusap1*, *ube2c*, *birc5a*, *birc5b*, *tpx2*, *top2a*, *ndc80*, *cks2*, *nuf2*, *cks1b*, *mki67*, *tmpoa*, *tmpob*, *cenpf*, *tacc3*, *smc4*, *ccnb2*, *ckap2l*, *aurkb*, *bub1*, *kif11*, *anp32e*, *tubb4b*, *gtse1*, *kif20ba*, *kif20bb*, *cdc20*, *ttk*, *kif2c*, *rangap1a*, *rangap1b*, *ncapd2*, *dlgap5*, *cdca8*, *ect2*, *kif23*, *hmmr*, *aurka*, *anln*, *lbr*, *ckap5*, *cenpe*, *ctcf*, *nek2*, *g2e3*, *gas2l3*, *cbx5* |

**Table S5. Model quality metrics.** Computed with COBRApy for the distributed reconstruction, the reference network used in this study, and two representative extracted models. Internal reactions exclude exchange, demand, sink and biomass reactions; a reaction is mass balanced if every element balances and charge balanced if charge balances. Annotation coverage counts reactions or metabolites with at least one database cross-reference.

| Metric | Zebrafish-GEM (distributed) | Reference network | FASTCORE default, ISV_tip | FASTCORE default, aDA |
|---|---|---|---|---|
| Reactions | 12,833 | 9,480 | 6,863 | 7,160 |
| Metabolites | 8,435 | 5,578 | 4,638 | 4,843 |
| Genes | 2,709 | 2,170 | 1,919 | 1,941 |
| Internal reactions | 11,165 | 8,466 | 5,852 | 6,149 |
| Mass balanced (%) | 99.2 | 99.5 | 99.5 | 99.5 |
| Charge balanced (%) | 97.9 | 98.9 | 99.0 | 99.1 |
| Metabolites with formula (%) | 99.9 | 99.9 | 99.9 | 99.9 |
| Metabolites with charge (%) | 100.0 | 100.0 | 100.0 | 100.0 |
| Reactions with GPR (%) | 70.7 | 73.6 | 79.0 | 79.9 |
| Reactions with annotation (%) | 100.0 | 100.0 | 100.0 | 100.0 |
| Metabolites with annotation (%) | 100.0 | 100.0 | 100.0 | 100.0 |
| Flux-consistent reactions, all exchanges open | 11,356 | — | — | — |
| Flux-consistent reactions, embryonic medium | 9,480 | 9,480 | — | — |

#### Supplementary Data S1

The workbook FluxSprout_Supplementary_Data.xlsx contains Tables S1–S5 in machine-readable form and the following data sheets, each generated by the analysis notebook.

| Sheet | Content |
|---|---|
| D1 | Genes differentially expressed between ISV_cyc and ISV_tip, with correlation to the cell-cycle scores (Fig. S2A) |
| D2 | Pathway marker scores, ISV_cyc minus ISV_tip, raw and adjusted for phase and depth (Fig. 2D) |
| D3 | Marker-level tests of the eight claims, raw and adjusted (Fig. 3, Table 5) |
| D4 | Flux-level tests between metacell models at U = 50 and U = 100 (Fig. 5D, Table 5) |
| D5 | Flux read-outs, loop gap and number of bound reactions for every metacell model (Fig. 5C–E) |
| D6 | E-Flux capacity sweep for every pseudobulk group model (Fig. 5A, Fig. S3) |
| D7 | Growth and limiting nutrient for each biomass objective on the embryonic medium |
| D8 | Oxidative phosphorylation factorial: objective, medium, maintenance demand, reaction set and capacity (Fig. 4B) |
| D9 | Loop-capable reactions with subsystem and GPR rule (Fig. 4D–E) |
| D10 | Knockout screen: growth inhibition of every screened gene in every group, one sheet per ensemble member (Fig. 6A, Table 4) |
| D11 | Robustness of every called gene across members and rules (Table 4) |
| D12 | Reactions that restore growth for each all-lineage call (Section 3.7, Fig. 6D) |
| D13 | Knockout inhibition across media for the FASTCORE default and GIMME members (Fig. 6C) |
| D14 | Published phenotypes recomputed with loop correction (Table 6) |

##### Code and data

All analyses are contained in a single Jupyter notebook that regenerates every figure, table and supplementary item from the two GEO matrices (GSM6138416, GSM6138417) and the distributed Zebrafish-GEM, with random seeds fixed. The reference network and all 108 context-specific models (27 groups in each of the four ensemble members) are provided as SBML Level 3 files that can be read directly with COBRApy, together with the notebook, the analysis code and every derived table. Software: Python 3.13, Scanpy 1.12, AnnData 0.13, leidenalg 0.12, scikit-learn 1.9, statsmodels 0.15, COBRApy 0.32, highspy 1.15 and matplotlib 3.11.
